# PXN Unlocks the Power of Public Gene Expression Data Through Cross-Technology Integration

**DOI:** 10.64898/2026.05.11.724309

**Authors:** Zhining Sui, Disa Yu, Arslan Erdengasileng, Jinfeng Zhang, Xing Qiu

## Abstract

The immense value of public gene expression repositories is constrained by the lack of compatibility among datasets generated from diverse experimental technologies. Differences in measurement scales, probe chemistries, and signal distributions create systematic discrepancies across platforms and laboratories. These inconsistencies make large-scale integrative analysis nearly impossible, even though such studies could achieve great statistical power and improved reproducibility. We introduce PXN, a probabilistic machine learning framework that captures a unified representation of biological signal across multiple gene expression technologies. Once trained, PXN can seamlessly translate data between multiple platforms, preserving informative biological variation while removing technology-specific biases. In benchmarking studies, PXN consistently outperforms existing normalization methods in cross-platform accuracy and substantially enhances the power of differential expression analysis. Importantly, we show that PXN is powerful enough to bridge even the most challenging technological divide—between microarray and RNA-seq. This capability provides a scalable route for integrating legacy microarray data with modern RNA-seq studies. By enabling direct comparison and integration of heterogeneous datasets, PXN unlocks the full potential of public repositories for future biological discovery and therapeutic innovation.

## Introduction

Over the past two decades, transcriptomic profiling has transformed biomedical research. Thousands of studies across diseases, model systems, and experimental conditions have collectively generated millions of gene-expression measurements, archived in public repositories such as the Gene Expression Omnibus (GEO; (Edgar, Domrachev et al. 2002)), ArrayExpress (Brazma, Parkinson et al. 2003), and The Cancer Genome Atlas (TCGA; (Cancer Genome Atlas Research, Weinstein et al. 2013)). These data represent an extraordinary scientific resource: a record of cellular states and disease phenotypes that could in principle be re-analyzed jointly to achieve better power and reproducibility (Hong and Breitling 2008), potentially leading to new biology (Sielemann, Hafner et al. 2020). Yet this vast wealth of information remains largely untapped (Moretto, Sonego et al. 2019). The challenge lies primarily in incompatibility among measurement technologies (Consortium 2014, Zhang, Shao et al. 2020). Each generation of transcriptomic assays—from early microarrays (Schena, Shalon et al. 1995) to modern RNA sequencing (RNA-seq; (Nagalakshmi, Wang et al. 2008, Wang, Gerstein et al. 2009)) has used distinct probe designs, signal detection chemistries, and dynamic ranges. These differences introduce technology-specific biases that lead to systematic differences in statistical properties across different platforms, such as mean expression levels, variances, and gene-gene correlations (Zhang, Shao et al. 2020, Borisov, Sorokin et al. 2022). As a result, data collected in separate studies cannot be directly compared, even when they examine the same tissue or biological process (Zhang, Shao et al. 2020). Among these divides, the gap between microarray and RNA-seq is the most striking: they differ not only in scale and noise characteristics, but also in how expression values are defined, normalized, and interpreted.

The consequence is a fragmented landscape of genomic knowledge. Instead of a single, unified resource, the research community is left with thousands of isolated datasets, each confined to its own technological context. This fragmentation limits the reproducibility of individual findings (Irizarry, Warren et al. 2005, Consortium 2006, Consortium 2014), reduces the statistical power of meta-analyses (Tseng, Ghosh et al. 2012, Qiu, Wu et al. 2013, Qiu, Hu et al. 2014), and prevents direct integration of legacy data with new studies (Zhao, Fung-Leung et al. 2014, Zhang, Shao et al. 2020). In effect, decades of valuable measurements are locked behind technical barriers, constraining our ability to build comprehensive, data-driven models of biology.

- **Biological integrity:** A common limitation of existing cross-platform normalization methods is that they do not leverage the gene–gene correlation structure that underlies coordinated biological regulation (Ideker, Thorsson et al. 2001, Li 2002). Prior work from our group and others has shown that inter-gene dependence strongly influences normalization (Qiu, Brooks et al. 2005,Qiu, Klebanov et al. 2005) and differential-expression analysis (Almudevar, Klebanov et al. 2006, Efron 2007, de la Fuente 2010, Liu, Zhang et al. 2017, Cui, Liu et al. 2021). Empirical evidence shows that these dependencies are largely driven by low-rank latent factors (Alter, Brown et al. 2000, Li 2002, Leek and Storey 2007). By leveraging probabilistic PCA (PPCA; (Tipping and Bishop 1999)), PXN captures this low-rank covariance structure in a form called “shared information” (SI, see *Supplementary Information* Section S3). This approach preserves gene-gene correlations, improves missing-value estimation, thereby enabling more accurate cross-platform normalization and increases the statistical power of downstream differential gene expression analyses (DGEA).
- **Computational efficiency and scalability:** PXN supports simultaneous normalization across more than two datasets, enabling seamless harmonization of multi-platform studies within a single unified model. PXN leverages probabilistic principal-component analysis to learn the shared covariance structure of biological variation directly from all datasets, allowing unbiased alignment between diverse technologies. It embraces a “*learn once, apply everywhere*” architecture: a single unified model trained from all datasets can translate expression profiles between any supported platform pair in either direction without retraining. Our estimation algorithm relies on conditional maximum likelihood and exploits Kronecker-structured matrix operations, singular value decomposition, and the Woodbury identity to avoid large matrix inversions and maintain numerical stability, making it highly scalable for large-scale high-throughput data.
- **Zero-shot model bridging:** PXN can perform “zero-shot” normalization (Kedzierska, Crawford et al. 2023, Cui, Wang et al. 2024) between disconnected platforms that lack direct paired samples in the training data. By mathematically synthesizing independently trained component models into a unified global network via their shared latent structure, PXN can accurately impute unobserved platform mappings. This is impossible or very difficult to implement with other standard pairwise approaches like Shambhala or MatchMixeR.

Together, these properties make PXN a general framework for cross-technology integration that offers reference-free probabilistic alignment across platforms and enables coherent, biologically consistent representations of transcriptomic landscapes. Through comprehensive simulations and real-world applications, we show that PXN unifies data once divided by technical boundaries, helping researchers to access the full value of public genomic repositories.

## Results

### Overview of the PXN Framework

Unlike traditional normalization methods that align datasets through pairwise platform transformations or empirical batch correction, PXN adopts a generative approach based on the PPCA to integrate gene expression data measured by multiple technological platforms. PXN models all observed datasets as realizations of a common latent biological signal that undergoes platform- and gene-specific transformations: offset, scaling, and noise addition. Each platform provides a distinct “view” of the same underlying transcriptomic landscape, rendered by its own measurement properties. By learning these relationships *jointly across all platforms*, PXN infers the shared low-dimensional structure that best explains the biological variation present in every dataset.

This modeling framework offers several key advantages. First, PXN explicitly models both gene- and platform-specific variability, which allows it to integrate heterogeneous datasets that differ not only in mean expression levels but also in measurement precision. Second, PXN can incorporate covariates such as age, sex, and clinical conditions into the model and adjust their confounding effects during normalization rather than as a post-hoc correction. Third, the probabilistic formulation enables us to derive principled parameter estimation and normalization procedures that attain theoretical optimality under mild assumptions. Specifically, the parameter estimation for PXN is carried out using an algorithm inspired by both the expectation-maximization (EM) algorithm and gradient descend (GD), which partitions parameters into manageable subsets and iteratively maximizes the conditional likelihood (details provided in *Methods* and *Supplementary Information*). Once trained, PXN performs cross-platform normalization using the empirical best linear unbiased predictor (EBLUP): given data from one technology, it predicts the expected expression values as they would appear on another platform. This operation enables *bi-directional translation* between datasets, allowing users to compare or combine studies measured by different technologies.

Importantly, PXN is designed for *multi-platform integration*. Rather than aligning data in pairs, it can learn a unified model across all available platforms, thereby avoiding inconsistencies that arise from sequential pairwise mappings. This global approach ensures that the learned latent structure captures the shared biological space underlying all platforms simultaneously.

Together, these features make PXN a scalable, reference-free, and biologically grounded solution for harmonizing large transcriptomic collections. Technical details of the PXN model and algorithms are described in the *Methods* section and *Supplementary Information*.

### Simulated Datasets

To evaluate the accuracy of PXN in harmonizing gene-expression data across technologies, we conducted a controlled simulation using measurements from various technical platforms. This simulated dataset consisted of 1,000 gene expression profiles across *N* = 240 subjects sequenced on six technical platforms, resulting in a 6,000×240 matrix (**Figure 1B**). From this full dataset, we extracted three training subsets: 1. Subset AB, a 2,000×60 matrix corresponding to cells A1 and B1 in **Figure 1B**, representing *n*_1_ = 60 subjects measured on platforms A and B. 2. Subset BCD, a 3,000×80 matrix (cells B2, C2, and D2) consisting of *n*_2_ = 80 subjects measured on platforms B, C, and D. 3. Subset EF, a 2,000×100 matrix (cells E3 and F3) composed of *n*_3_ = 100 subjects measured on platforms E and F.

**Figure 1.**
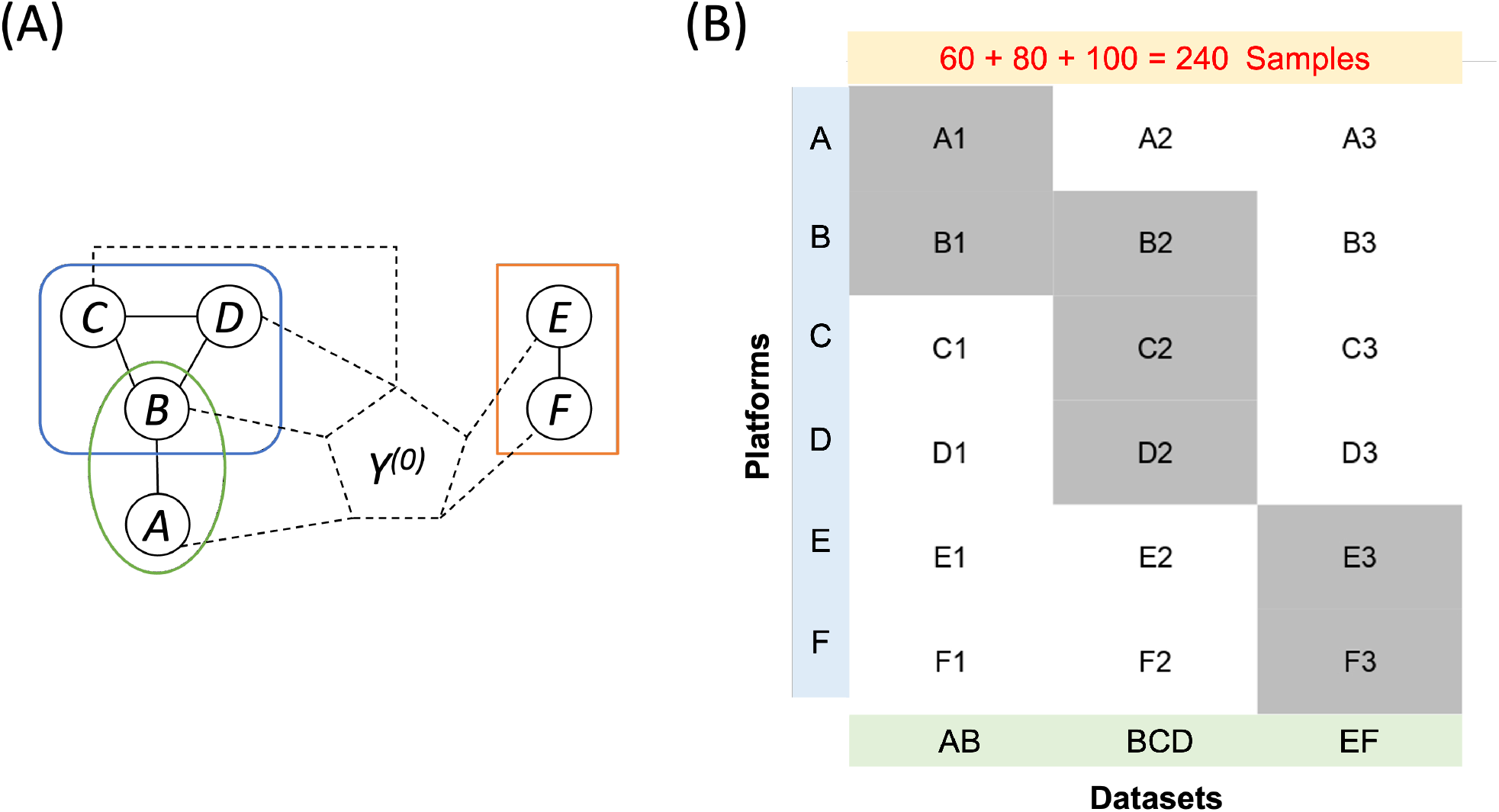
Graphical illustration of the simulated datasets. (A): The pairing structures of six sets of simulated data. PXN can be applied separately to the green region (A and B, with *n*_1_ = 60 paired samples), blue region (B, C, and D with *n*_2_ = 80 paired samples), or the orange region (E and F, with *n*_3_ = 100 paired samples). The results from these analyses can be bridged via internal connections (shown as broken lines) between these platforms to the latent platform represented by *Y*^(0)^ in the PXN model. (B) An alternative visualization of this pairing structure in terms of three subsets of paired data: AB (*n*_1_ = 60), BCD (*n*_2_ = 80), and EF (*n*_3_ = 100). Shaded cells are the observed data and unshaded ones represent missing platforms.

The remaining measurements (white cells in **Figure 1B**, e.g., A2, B3) corresponded to unobserved platforms and were excluded from the training data. They were retained as ground-truth data for evaluating the accuracy of cross-platform normalization procedures. The overall structure of these simulated data is illustrated in **Figure 1**, and technical details of data generation are provided in *Supplementary Information* Section S5.

### Simulation Analysis I: Accuracy of Cross-platform Normalization

We compared PXN with two representative normalization frameworks: MatchMixeR and Shambhala-2 in Simulation Analysis I. We also examined how specific design components affect PXN’s performance. Four PXN variants were tested: the full model (**PXN**, including covariate adjustment and gradient-descent refinement), **PXN-noX** (without covariate adjustment), **PXN-noGD** (without gradient-descent refinement), and **PXN-noX-noGD** (without both).

Ten-fold cross-validation was used for model training and evaluation: 90% of samples were used for training, and the remaining 10% were used for prediction. Each trained model was applied to infer gene-expression levels on platform B from the corresponding platform A measurements. Prediction accuracy was quantified using the mean squared prediction error (MSPE). Shambhala-2, which applies direct sample-wise transformation without training, was evaluated by its residual sum of squares.

We first examined how PXN’s performance varies with different choices of latent dimensionality (*L*) and two modeling components (with or without covariate adjustment and gradient descent refinement) using dataset AB (*n* = 60). The results are shown in **Figure 2**. For comparison, results from two MatchMixeR implementations (MM and MM-OLS) are displayed in **Figure 2** as red and blue horizontal reference lines. Across all values of *L*, every PXN variant achieved a lower prediction error than both MatchMixeR baselines. The full PXN model (with both covariate adjustment and gradient-descent refinement) consistently produced the smallest MSPE. Furthermore, the optimal latent dimension (identified through an independent and computationally efficient five-fold cross-validation using PXN-noGD) was stable across all four variants (*L*^*^ = 7) and largely consistent with the observed minimum MSPE.

**Figure 2.**
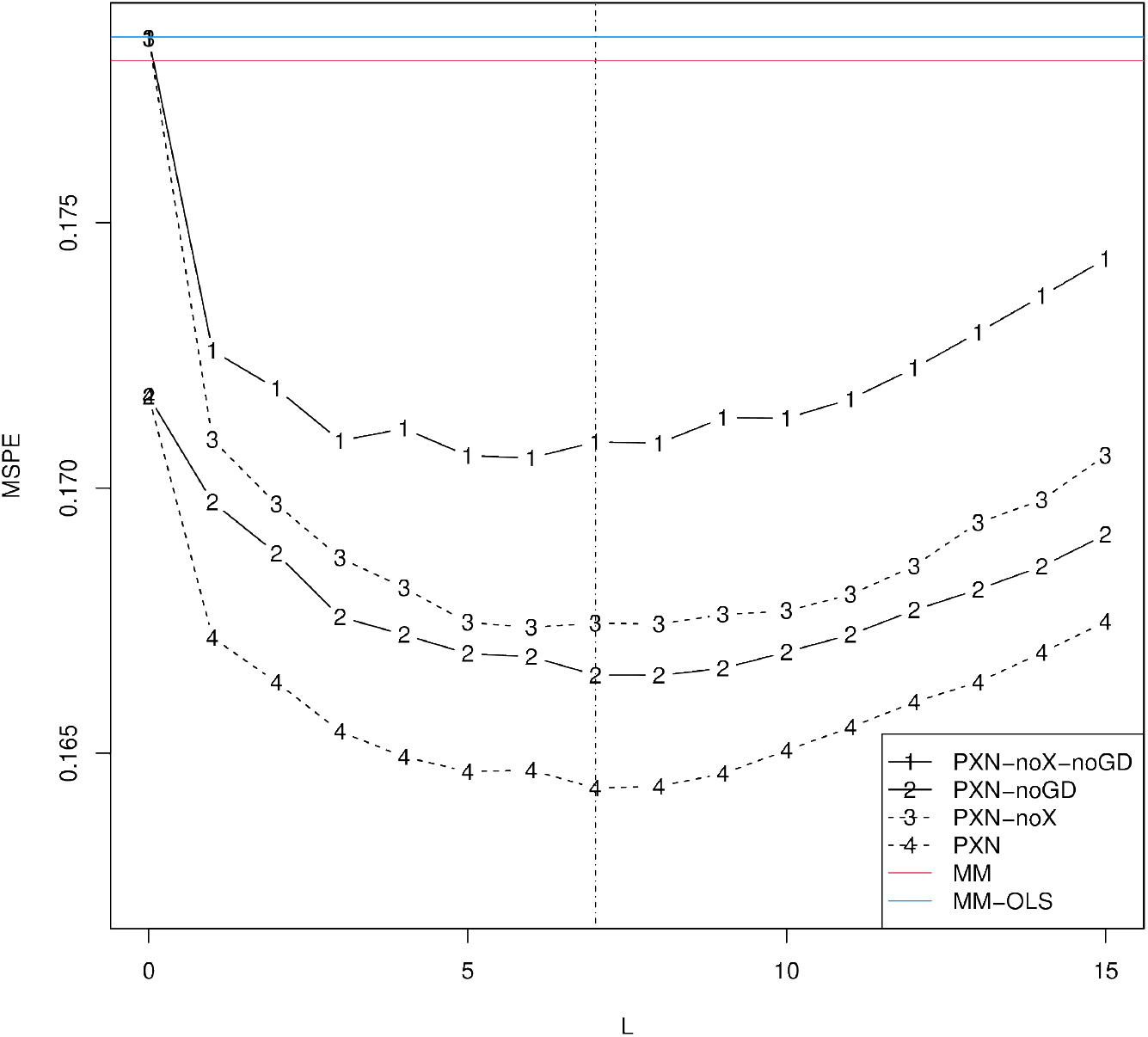
Accuracy of cross-platform normalization across PXN modeling configurations. Simulated dataset AB (*n* = 60) was used in this comparison. The red and blue horizontal reference lines represent MSPEs of MM and MM-OLS (two variants of MatchMixeR). Curves represent MSPEs of four variants of PXN. The dashed vertical line represents the optimal *L*^*^ selected by an independent five-fold cross-validation procedure using PXN-noGD.

We next extended the evaluation to multiple normalization scenarios using both the AB and BCD datasets, covering diverse combinations of source and target platforms. Across all four prediction tasks, PXN consistently achieved the lowest MSPE, outperforming both MatchMixeR variants and Shambhala-2 by substantial margins (**Table 1**). Notably, these improvements were observed regardless of the source– target direction, underscoring the symmetry and stability of PXN’s multi-platform normalization.

**Table 1.** Accuracy of Cross-platform Normalization. Mean squared prediction errors (MSPEs) from 10-fold cross-validation for cross-platform normalization tasks using the AB and BCD datasets. Each entry represents the MSPE when predicting gene expression on the target platform (right of arrow) from the source platform (left of arrow). The latent dimension *L* in PXN was selected via an independent five-fold cross-validation using PXN-noGD. The “Original” row corresponds to a naïve baseline that uses unnormalized target platform data as predictions.

| | A1 → B1 | B2 → C2 | B2 → D2 | C2 → D2 |
| --- | --- | --- | --- | --- |
| Original | 0.9658 | 0.9903 | 1.7351 | 0.9667 |
| PXN | 0.1643 | 0.1344 | 0.1881 | 0.1885 |
| MM | 0.1781 | 0.1481 | 0.2064 | 0.2052 |
| MM-OLS | 0.1785 | 0.1483 | 0.2067 | 0.2055 |
| Shambhala-2 | 2.0748 | 1.2342 | 1.6335 | 2.1554 |

### Simulation Analysis II: Multi-platform Model Bridging

A common barrier to cross-study integration is the lack of directly paired samples between certain platforms. PXN addresses this issue via **model bridging**, a unique technique that synthesizes a unified global PXN model from several smaller, independently trained models by leveraging the shared latent structure (*Methods* and *Supplementary Information* Section S4). In this simulation, we bridged the three independently trained subsets into a global PXN model, using the rounded weighted average of the optimal values *L*^*^ from each model as the global latent dimensionality.

We compared predictions from the bridged model with those from individually trained PXN models, as well as with unnormalized baselines (“Original”). Notably, we evaluated PXN’s ability to normalize between platforms that have *no direct pairing* in the training data, e.g., between platforms A and C (connected only through B) and between A and F (completely disconnected). This type of harmonization is known as zero-shot normalization, which is very challenging in general and impossible for MatchMixeR and individually trained PXN models without bridging.

For completeness, we also included a “No pairing” baseline model. In this approach, we deliberately ignore the 1-to-1 pairing structure in the training data. PXN parameters are estimated independently for each platform as if these datasets were completely disjoint. The normalization mappings (e.g., A → B and A → C) are then derived from bridging these independently fitted PXN models. We use this method to demonstrate that jointly modeling the shared latent space across paired samples is advantageous.

As shown in **Table 2**, explicitly modeling the pairing structure yields substantial improvements. Both the direct and the globally bridged PXN models dramatically reduced prediction errors when compared to models that ignored the paired design (“No pairing”), which in turn were substantially better than unnormalized baselines (“Original”).

**Table 2.** Accuracy of Cross-platform Normalization. Mean squared prediction errors (MSPEs) from 10-fold cross-validation for various cross-platform normalization tasks using PXN. “Original” refers to unnormalized target values. “Direct” indicates PXN models trained on pairing groups AB, BCD, and EF separately. “Bridged” refers to the overall model that combines all three direct PXN models. “No pairing” corresponds to a bridged model trained without leveraging platform pairings. Note that predictions for A2 → C2 and A3 → F3 are only possible using bridged models.

| | A1 → B1 | A2 → C2 | B2 → C2 | A3 → F3 | E3 → F3 |
| --- | --- | --- | --- | --- | --- |
| Original | 0.9658 | 1.7313 | 0.9903 | 7.1531 | 1.0988 |
| Direct | 0.1672 | N/A | 0.1356 | N/A | 0.2161 |
| Bridged | 0.1597 | 0.1425 | 0.1313 | 0.2354 | 0.2093 |
| No pairing | 0.2589 | 0.2367 | 0.2330 | 0.3840 | 0.6043 |

Importantly, the bridged model successfully performed zero-shot normalization for disconnected platforms (A2 → C2 and A3 → F3) with MSPEs (0.1425 and 0.2354, respectively) that are on par with connected cases, e.g., A1 → B1 (0.1597) and E3 → F3 (0.2093). Furthermore, for cases where direct pairing data were available (A1 → B1, B2 → C2, and E3 → F3), the bridged model performed on par with or even slightly better than the directly targeted individual models.

It is worth noting that PXN achieves this multi-platform capability **without retraining**: a single globally bridged model was able to translate expression profiles across *any combination* of supported technologies. This makes PXN particularly attractive for harmonizing large transcriptomic repositories, where constructing all pairwise normalization models would be impractical. Moreover, because PXN learns a shared latent space across all platforms, it avoids the inconsistencies that can arise from sequential or chain-based mappings.

Together, these results demonstrate that the multi-platform training framework of PXN can effectively leverage shared information across technologies, providing a coherent and scalable solution for cross-technology harmonization, especially in settings where direct platform pairings are sparse or incomplete.

### Simulation Analysis III: Impact on Differential Expression Analysis

We next assessed how cross-platform normalization affects downstream gene–differential-expression analysis (GDEA). Two complementary scenarios were considered.

### Sub-analysis IIIa: Balanced covariates and paired samples

To determine whether normalization introduces statistical bias that inflates type I error or reduces power in DGEA, we first evaluated a perfectly balanced scenario. Platform A1 samples were normalized to Platform B (denoted by 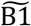) by PXN, Shambhala-2, MatchMixeR, or an unnormalized baseline (“Original”). They were combined with their paired B1 counterparts to form dataset (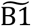, B1). Because samples in A1 and B1 represent the exact same *n* = 60 subjects sequenced on platforms A and B, they share identical values for all three simulated covariates: Treatment, Age, and an additional confounder *x* (see section S5, *Supplementary Information*). Consequently, covariates are perfectly balanced across the two platforms, making normalization unnecessary in theory.

To identify differentially expressed genes (DEGs) associated with Treatment and Age, we evaluated three statistical approaches:

1. **OLS**: A multiple regression model incorporating all three covariates was applied directly to the combined dataset (B1, 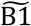). DEGs were selected based on the standard regression t-test for Treatment and Age.
2. **LMER**: A linear mixed-effects model was applied to the combined dataset (B1, 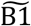). It extends the OLS approach by adding a subject-specific random intercept to account for the pairing structure between B1 and 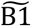. DEGs were selected by regression t-test with Satterthwaite approximation for degrees of freedom.
3. **pcomb**: The OLS model was applied to B1 and 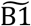 independently. The covariate-specific p-values from the two datasets were then combined using Fisher’s method.

We evaluated power, false-positive rate (FPR), and area under the curve (AUC) over 100 repetitions. As shown in **Table 3**, only LMER appropriately controlled the false-positive rate (FPR), whereas OLS and pcomb failed by ignoring sample pairing. Furthermore, normalization had minimal impact in this perfectly balanced setting: the best AUC was attained with unnormalized data. This confirms that when covariates are perfectly aligned across platforms, cross-platform normalization is unnecessary, provided that the paired data structure is modeled to ensure valid inference.

**Table 3.** Summary statistics of Simulation Analysis IIIa. All results were averaged from 100 repetitions and presented on a percentage scale for better readability. Highlighted are the best AUC statistics among all methods.

|  | Original |  |  | PXN |  |  | Shambhala-2 |  |  | MatchMixeR |  |  |
| --- | --- | --- | --- | --- | --- | --- | --- | --- | --- | --- | --- | --- |
|  | OLS | LMER | pcomb | OLS | LMER | pcomb | OLS | LMER | pcomb | OLS | LMER | pcomb |
| **Power** |  |  |  |  |  |  |  |  |  |  |  |  |
| Treatment | 80.7 | 71.6 | 78.0 | 85.2 | 72.8 | 78.9 | 80.1 | 72.8 | 79.3 | 84.9 | 72.3 | 78.3 |
| Age | 81.7 | 77.4 | 80.3 | 83.0 | 76.6 | 79.3 | 80.2 | 77.3 | 79.6 | 83.4 | 77.5 | 79.8 |
| **FPR** |  |  |  |  |  |  |  |  |  |  |  |  |
| Treatment | 6.3 | 2.2 | 4.8 | 13.3 | 4.2 | 8.2 | 5.5 | 3.0 | 5.8 | 10.6 | 2.8 | 5.1 |
| Age | 7.7 | 2.8 | 5.9 | 12.9 | 3.5 | 6.7 | 6.9 | 3.7 | 7.4 | 11.3 | 3.0 | 5.4 |
| **AUC** |  |  |  |  |  |  |  |  |  |  |  |  |
| Treatment | 94.2 | 94.0 | 93.8 | 93.1 | 92.7 | 92.2 | 94.1 | 93.8 | 93.6 | 93.7 | 93.5 | 93.5 |
| Age | 91.1 | 91.1 | 91.2 | 90.6 | 90.5 | 90.4 | 90.9 | 90.9 | 90.7 | 91.0 | 90.9 | 90.9 |

### Sub-analysis IIIb: Unbalanced covariates and unpaired samples

The second scenario examined a more realistic case in which covariates differed substantially across platforms. We trained a globally bridged PXN model using all simulated paired datasets (AB, BCD, and EF), and used it to normalize an independent dataset A4 (*n* = 30) to platform B (denoted as 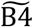). These normalized samples were then combined with dataset B5 (*n* = 50). Unlike the previous perfectly paired scenario, the Treatment and Age distributions were intentionally made unbalanced across the two platforms to emulate confounding patterns that often complicate real-world integrative analysis. Because no pairing samples were presented in this dataset, the OLS and pcomb approaches were employed to identify DEGs.

As summarized in **Table 4**, applying the OLS model to unnormalized data completely failed to control type I error due to the confounding effects introduced by the cross-platform covariate imbalance. While the pcomb method successfully controlled the false-positive rate (FPR) by performing platform-specific inferences, it severely sacrificed statistical power. In comparison, PXN-based approaches produced the strongest overall performance. They had tight type I error control with substantially higher power and Area Under the Curve (AUC) for both Treatment and Age, especially when used with the OLS approach.

**Table 4.** Summary statistics of Simulation Analysis IIIb. All results are averaged from 100 repetitions and presented in percentage scale for better readability. Highlighted are the best AUC statistics among all methods.

|  | Original |  | PXN |  | Shambhala-2 |  | MatchMixeR |  |
| --- | --- | --- | --- | --- | --- | --- | --- | --- |
|  | OLS | pcomb | OLS | pcomb | OLS | pcomb | OLS | pcomb |
| **Power** |  |  |  |  |  |  |  |  |
| Treatment | 65.0 | 71.2 | 90.4 | 76.2 | 65.1 | 64.4 | 86.2 | 70.6 |
| Age | 76.5 | 66.0 | 77.4 | 68.1 | 67.5 | 67.6 | 76.9 | 66.0 |
| **FPR** |  |  |  |  |  |  |  |  |
| Treatment | 20.8 | 2.9 | 3.0 | 3.0 | 9.7 | 7.6 | 3.3 | 2.9 |
| Age | 30.9 | 2.5 | 4.3 | 2.1 | 2.8 | 5.5 | 4.8 | 2.4 |
| **AUC** |  |  |  |  |  |  |  |  |
| Treatment | 78.3 | 94.5 | 97.7 | 95.8 | 83.8 | 86.8 | 96.5 | 94.2 |
| Age | 82.9 | 87.6 | 89.4 | 88.4 | 88.4 | 86.4 | 89.1 | 87.4 |

The superiority of the unified OLS model over the platform-specific pcomb approach on PXN-harmonized data highlights a fundamental advantage of our method. The pcomb strategy is inherently a conservative, “divide-and-conquer” approach: it protects against inflated type I errors when platform discrepancies remain after normalization, but it sacrifices statistical efficiency in doing so. The fact that OLS achieved the highest power without inflating the FPR demonstrates that PXN aligned 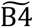 and B5 almost perfectly. By transforming disparate datasets into true homogenous counterparts, PXN eliminates the need for conservative statistical guardrails and allows researchers to achieve the maximum statistical power of a pooled dataset.

### Real-data Demonstration I: Accuracy of Pairwise Cross-platform Normalization

We evaluated PXN on the CellMiner molecular profiling resource from the U.S. National Cancer Institute, which provides multi-platform transcriptomic measurements for the NCI-60 panel of human cancer cell lines. This dataset consists of data generated on six technical platforms. Specifically, it includes four generations of Affymetrix GeneChip platforms: Human Genome U95 (HG-U95 A–E; five-chip set; ~65,000 probe sets), Human Genome U133 (HG-U133 A–B; two-chip set; ~44,000 probe sets), Human Genome U133 Plus 2.0 (~47,000 transcripts), and the exon-level Human Exon 1.0 ST array. In addition, CellMiner provides an Agilent Whole Human Genome Microarray (4×44K) and an RNA-seq dataset generated on the Illumina HiSeq 2000 platform. All data were processed using standard summarization pipelines: the Affymetrix arrays were summarized using GCRMA, RMA, or MAS5; the Agilent data were processed via GeneSpring; and the RNA-seq reads were aligned to the hg19 reference genome using STAR with accompanying quality-control summaries. For readability, original CellMiner platform names were mapped to short labels (A = HG-U133, B = HG-U133 Plus 2.0, C = HG-U95, D = HuEx-1.0-ST, E = Agilent, F = RNA-seq).

Across the six expression platforms, 57 matched samples were observed in common. We assessed cross-platform predictive accuracy using 10-fold cross-validation with a single set of fold assignments reused across all platforms and models. In each fold, PXN models and two MatchMixeR variants were trained on 90% of paired samples and evaluated on the held-out 10% by bidirectional prediction. We identified a subset of 7,441 genes measured across all six platforms. This common gene set serves as the primary basis for direct gene-level cross-platform comparisons. For each trained model and each ordered source-target platform pair it supports, we predicted target-platform expression from the source platform and computed the mean squared prediction error (MSPE) against the observed target expression in the same held-out samples. MSPE was computed on the gene set used to fit the corresponding model, ensuring that comparisons were made within a consistent feature space. For all six platforms, our evaluation produced a total of 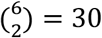 ordered pairwise normalization tasks. The results of these cross-validated predictions are summarized in **Table 5**.

**Table 5.**
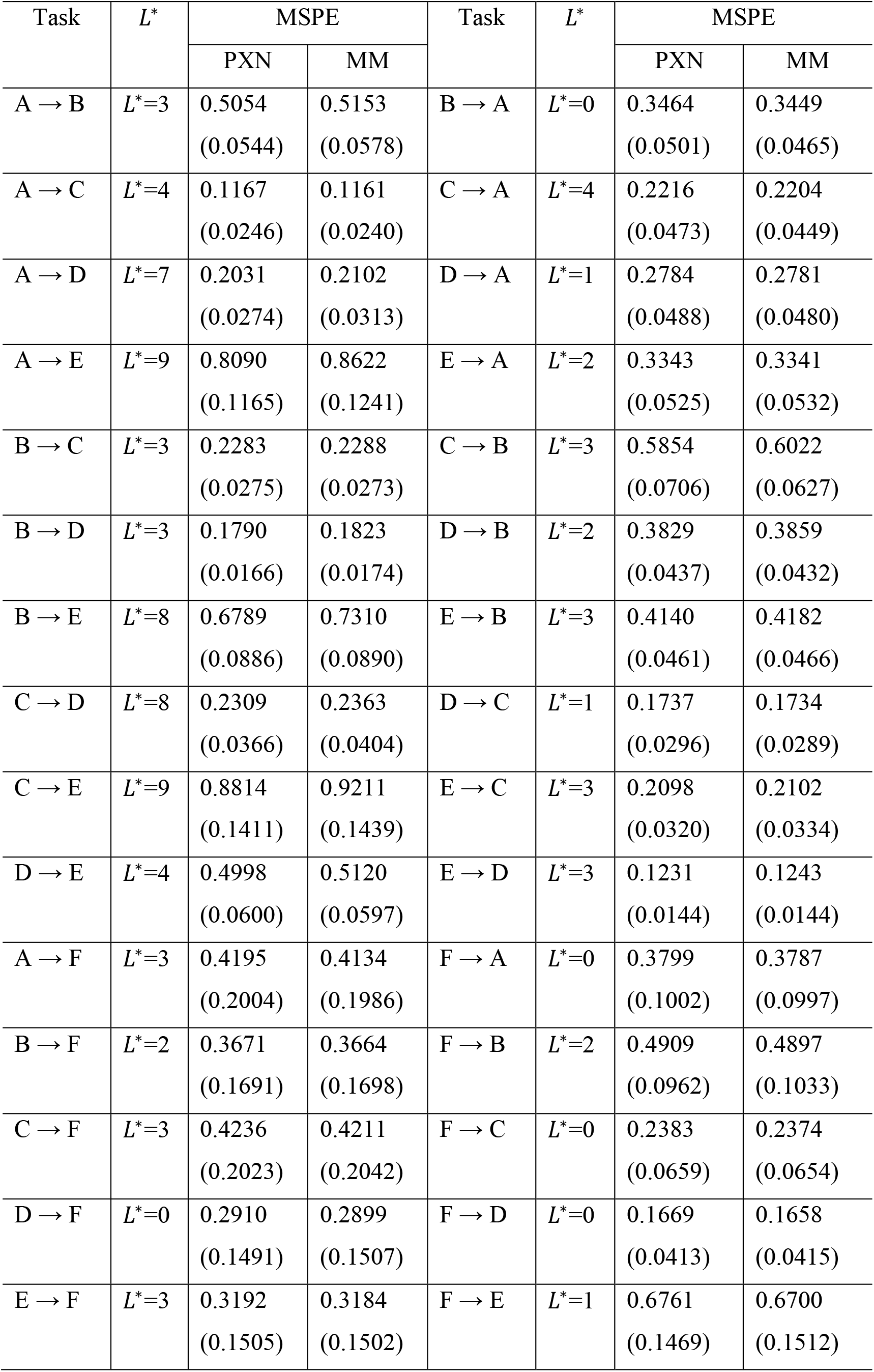
Pairwise normalization accuracy for Real-data Demonstration I. For each ordered platform-to-platform normalization task, we report the latent dimension *L*^*^ selected for PXN (minimizing PXN MSPE) and the corresponding MSPE (mean ± SD across cross-validation folds) for PXN and MM. The mean MSPE for all 30 tasks is **0.3701** for PXN and **0.3803** for MatchMixeR,

| Task | $L^*$ | MSPE | | Task | $L^*$ | MSPE | |
| --- | --- | --- | --- | --- | --- | --- | --- |
|  |  | PXN | MM |  |  | PXN | MM |
| $A \rightarrow B$ | $L^*=3$ | 0.5054<br>(0.0544) | 0.5153<br>(0.0578) | $B \rightarrow A$ | $L^*=0$ | 0.3464<br>(0.0501) | 0.3449<br>(0.0465) |
| $A \rightarrow C$ | $L^*=4$ | 0.1167<br>(0.0246) | 0.1161<br>(0.0240) | $C \rightarrow A$ | $L^*=4$ | 0.2216<br>(0.0473) | 0.2204<br>(0.0449) |
| $A \rightarrow D$ | $L^*=7$ | 0.2031<br>(0.0274) | 0.2102<br>(0.0313) | $D \rightarrow A$ | $L^*=1$ | 0.2784<br>(0.0488) | 0.2781<br>(0.0480) |
| $A \rightarrow E$ | $L^*=9$ | 0.8090<br>(0.1165) | 0.8622<br>(0.1241) | $E \rightarrow A$ | $L^*=2$ | 0.3343<br>(0.0525) | 0.3341<br>(0.0532) |
| $B \rightarrow C$ | $L^*=3$ | 0.2283<br>(0.0275) | 0.2288<br>(0.0273) | $C \rightarrow B$ | $L^*=3$ | 0.5854<br>(0.0706) | 0.6022<br>(0.0627) |
| $B \rightarrow D$ | $L^*=3$ | 0.1790<br>(0.0166) | 0.1823<br>(0.0174) | $D \rightarrow B$ | $L^*=2$ | 0.3829<br>(0.0437) | 0.3859<br>(0.0432) |
| $B \rightarrow E$ | $L^*=8$ | 0.6789<br>(0.0886) | 0.7310<br>(0.0890) | $E \rightarrow B$ | $L^*=3$ | 0.4140<br>(0.0461) | 0.4182<br>(0.0466) |
| $C \rightarrow D$ | $L^*=8$ | 0.2309<br>(0.0366) | 0.2363<br>(0.0404) | $D \rightarrow C$ | $L^*=1$ | 0.1737<br>(0.0296) | 0.1734<br>(0.0289) |
| $C \rightarrow E$ | $L^*=9$ | 0.8814<br>(0.1411) | 0.9211<br>(0.1439) | $E \rightarrow C$ | $L^*=3$ | 0.2098<br>(0.0320) | 0.2102<br>(0.0334) |
| $D \rightarrow E$ | $L^*=4$ | 0.4998<br>(0.0600) | 0.5120<br>(0.0597) | $E \rightarrow D$ | $L^*=3$ | 0.1231<br>(0.0144) | 0.1243<br>(0.0144) |
| $A \rightarrow F$ | $L^*=3$ | 0.4195<br>(0.2004) | 0.4134<br>(0.1986) | $F \rightarrow A$ | $L^*=0$ | 0.3799<br>(0.1002) | 0.3787<br>(0.0997) |
| $B \rightarrow F$ | $L^*=2$ | 0.3671<br>(0.1691) | 0.3664<br>(0.1698) | $F \rightarrow B$ | $L^*=2$ | 0.4909<br>(0.0962) | 0.4897<br>(0.1033) |
| $C \rightarrow F$ | $L^*=3$ | 0.4236<br>(0.2023) | 0.4211<br>(0.2042) | $F \rightarrow C$ | $L^*=0$ | 0.2383<br>(0.0659) | 0.2374<br>(0.0654) |
| $D \rightarrow F$ | $L^*=0$ | 0.2910<br>(0.1491) | 0.2899<br>(0.1507) | $F \rightarrow D$ | $L^*=0$ | 0.1669<br>(0.0413) | 0.1658<br>(0.0415) |
| $E \rightarrow F$ | $L^*=3$ | 0.3192<br>(0.1505) | 0.3184<br>(0.1502) | $F \rightarrow E$ | $L^*=1$ | 0.6761<br>(0.1469) | 0.6700<br>(0.1512) |

Overall, PXN achieved a lower global average MSPE (0.3701) compared to MatchMixeR (0.3803). Notably, the optimal latent dimension *L*^*^ selected for PXN was highly predictive of relative model performance (**Table 5**). When the data exhibited a low complexity latent space (*L*^*^ ∈ {0,1}), MatchMixeR was slightly but consistently better than PXN. Conversely, when the between-gene correlation structure supported a higher-dimensional shared latent structure (*L*^*^ ∈ {7,8,9}), PXN unequivocally outperformed MatchMixeR. This association suggests that *L*^*^ may serve as a practical, data-driven indicator for selecting the best normalization method for a given task.

### Real-data Demonstration II: Bridging Enables Normalization Beyond Directly Paired Models

In this real data demonstration, we evaluate the practical utility of model bridging on the CellMiner data, which consists of paired samples from six technical platforms. A practical question is how much predictive accuracy is lost when a normalization task must be performed **indirectly** through model bridging, rather than by training a dedicated pairwise model. We fit all 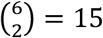 pairwise PXN models and then constructed all 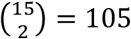 two-way bridged models. A pairwise model is denoted by the platforms used for training, such as [AB], which provides the direct baseline for normalizing between A and B. A bridged model is denoted by the combination of its component pairwise models, such as [AB, AC]. For a given directional normalization task, we defined a PXN model as *applicable* if the union of platforms covered by this model contains both the source and target platforms. For example, both the bridged model [AB, AC] and its first component model [AB] are applicable for tasks A → B, but only the bridged model is applicable for task B → C. Of course, either of them is applicable for task A→ D. Under this criterion, each directional task can be carried out using 30 applicable bridged models, providing multiple potential normalization routes.

To evaluate zero-shot bridging against the direct-pairing baseline, we categorized applicable bridged models by whether they include the direct source-target component. Using A → B as an example, 16 of 30 applicable bridged models do not include the direct-pairing component [AB], therefore we must normalize A → B via *zero-shot bridging*. The remaining 14 models include the [AB] component, which is deliberately bridged with an additional component (e.g., [BC] or [EF]). Those are the *perturbed* models designed to quantify the effect of injecting *extraneous structure* when a direct pairing already exists.

For each directional task, we averaged MSPE across applicable zero-shot bridged models (16 models per task) and across applicable perturbed models (14 models per task) and recorded the corresponding direct-pairing PXN baseline MSPE for that task. These detailed direction-level summaries are provided in Supplementary Table S3. Higher-level summaries of these results are provided in **Table 6**.

**Table 6.** Summary of Real-data Demonstration II: Task-averaged performance of bridged normalization. For each ordered platform-to-platform normalization task, we compare three approaches. “Direct” denotes the pairwise PXN model trained on the source–target platform pair. “Zero-Shot” denotes a bridged model that performs the same task without using the corresponding source–target pairwise component. “Perturbed” denotes a setting in which the direct model is intentionally bridged with an extraneous pairwise model involving unrelated platforms to assess the effect of introducing extra, potentially mismatched data when a direct mapping is already available. MSPE is computed by 10-fold cross-validation. Values are obtained by first averaging MSPE over applicable models within each task (as reported in Supplementary Table S3) and then averaging these task-level summaries across tasks (overall and stratified by whether the task involves Platform F). Ratios summarize relative error inflation of zero-shot bridging compared with direct and perturbed baselines.

|  | Average MSPE for Direct Models | Average MSPE for Bridged Models |  | Average MSPE Ratio |  |
| --- | --- | --- | --- | --- | --- |
|  |  | Zero-Shot | Perturbed | Zero-Shot / Direct | Perturbed / Direct |
| All tasks (30 tasks) | 0.3901 | 0.4050 | 0.3922 | 1.038 | 1.005 |
| Tasks not involving Platform F (20 tasks) | 0.3796 | 0.4012 | 0.3861 | 1.057 | 1.017 |
| Tasks involving Platform F (10 tasks) | 0.4112 | 0.4128 | 0.4044 | 1.004 | 0.983 |

Overall, PXN bridging provides a practical solution when the directly paired source-target model is unavailable. Across all 30 tasks, the average MSPE under zero-shot bridging was 0.4050, compared with 0.3901 for the pairwise baseline, corresponding to a 3.8% inflation. This modest inflation of MSPE reflects the anticipated trade-off of unmodeled heterogeneity in real-world data. A closer look revealed that, for tasks involving Platform F (Illumina HiSeq 2000, the only RNA-seq platform), the zero-shot bridged MSPE was essentially indistinguishable from the pairwise baseline. The slight performance penalty of zero-shot bridging was largely related to tasks harmonizing between legacy microarray platforms. Together, these results show that PXN’s bridging framework can support cross-platform normalization in settings where the directly paired source-target dataset is unavailable, while maintaining competitive prediction error, particularly for normalizations tasks involving the RNA-seq platform. This constitutes a key advantage over methods that rely strictly on direct pairing.

As a secondary analysis, we evaluated how much an otherwise well-specified direct normalization task could be affected when it is perturbed by bridging in additional platforms not required for the task. Across all tasks, the perturbed setting remained very close to the direct pairwise baseline (mean MSPE 0.3922 vs 0.3901; representing approximately 0.5% inflation). These results indicate that introducing additional, unnecessary platforms into the bridged model has negligible adverse impact on normalization accuracy.

In summary, when direct pairing is unavailable, PXN zero-shot bridging enables cross-platform normalization with only a modest loss in accuracy relative to directly trained pairwise models. When direct pairing is available, our perturbed setting demonstrates that the bridging mechanism remains highly robust to the injection of extraneous platforms.

### Real-data Demonstration III: Integrating Microarray and RNA-seq Data in DGEA

A realistic and challenging question for cross-technology analysis is whether harmonization with PXN (and other normalization methods) can improve statistical power while still maintaining proper type I error control. To address this, we applied DGEA to the TCGA dataset previously analyzed in the MatchMixeR study. This dataset includes 14,589 genes measured in 524 tumor and 58 normal samples (one sample excluded for quality reasons), with expression profiled on two paired technical platforms: Agilent G4502A (microarray) and Illumina HiSeq 2000 (a.k.a. GPL11154; RNA-seq). Because the true differentially expressed genes (DEGs) are unknown, we constructed a gold-standard DEG set by applying *limma* separately to each platform and selecting genes with consistent effect directions and sufficiently large effect sizes (see Section 6, *Supplementary Information*).

To evaluate how normalization affects downstream inference, we constructed training and testing splits consisting of small subsets of unpaired samples, mimicking real-world cross-platform integration scenarios. Two cases were considered, and the results are summarized in **Table 7**:

**Table 7.** Summary of Real-data Demonstration III. Expression data from Agilent Custom Gene Expression Microarray G4502A were normalized to align with the RNA-seq data from Illumina HiSeq 2000, GPL11154. All results were presented on a percentage scale for better readability. Highlighted are the best AUC statistics among all methods.

| Method | Balanced |  |  | Unbalanced |  |  |
| --- | --- | --- | --- | --- | --- | --- |
|  | Power | FPR | AUC | Power | FPR | AUC |
| Original | 6.55 | 0.01 | 89.82 | 92.97 | 92.76 | 56.22 |
| PXN ($L = 1$) | 72.74 | 2.90 | 93.55 | 76.96 | 3.35 | 94.12 |
| PXN ($L = 2$) | 74.10 | 3.06 | 93.75 | 76.89 | 3.37 | 94.08 |
| PXN ($L = 3$) | 74.34 | 3.16 | 93.75 | 76.94 | 3.37 | 94.10 |
| PXN ($L = 4$) | 74.79 | 3.22 | 93.86 | 76.96 | 3.36 | 94.11 |
| PXN ($L = 5$) | 74.96 | 3.27 | 93.88 | 76.96 | 3.35 | 94.11 |
| PXN ($L = 8$) | 75.44 | 3.37 | 93.98 | 76.96 | 3.34 | 94.11 |
| MatchMixeR | 72.81 | 2.98 | 93.50 | 76.38 | 3.39 | 93.94 |
| Shambhala-2 | 60.81 | 3.15 | 89.29 | 53.58 | 9.66 | 80.41 |

i. **Balanced case:** Tumor and normal samples were equally represented across platforms. Each test set included 8 tumor and 5 normal samples from each platform (*n* = 26 in total). The rest were used as the training set.
ii. **Unbalanced case:** Each test set comprised 10 microarray tumor samples, 15 RNA-seq tumor samples, and 10 RNA-seq normal samples (*n* = 35 in total). The rest were used as the training set. By omitting microarray normal samples, we introduced a platform-specific confounding effect.

Because the test data were unpaired, the LMER method was not applicable. Likewise, p-value combination (pcomb) was excluded from the unbalanced scenario, as the microarray platform lacked normal samples. In both cases, we performed DE analysis using the OLS approach and summarized the results across 100 training-test splits in **Table 7**.

As expected, combining unnormalized data (“Original”) yielded very poor results. In the balanced case, cross-platform technical variations completely overwhelmed the biological signal, resulting in near-zero statistical power (6.55%). In the unbalanced case, platform-specific sample composition severely confounded the inference, resulting in extremely high FPR (92.76%) and much reduced AUC (56.22%). All three normalization methods substantially improved downstream DGEA performance, but PXN consistently achieved the best overall performance. Shambhala-2 improved performance relative to “Original” but remained markedly inferior to PXN and MatchMixeR across all evaluation metrics, which is consistent with our simulation studies. PXN and MatchMixeR both achieved large power gains in the balanced design and successfully restored strict type I error control in the unbalanced design. However, for nearly every choice of latent dimension, PXN achieved better AUC values than MatchMixeR. Furthermore, MatchMixeR can only integrate two paired platforms at a time and requires a user-specified normalization direction, limiting its applicability in multi-platform settings.

Taken together, these results demonstrate that PXN reliably bridges the microarray–RNA-seq divide for downstream analysis even when sample composition differs significantly across platforms. By harmonizing expression profiles across technologies, PXN provides a robust foundation for combining legacy microarray data with modern RNA-seq studies, which could greatly expand the scope and statistical efficiency of integrative genomic analysis.

## Discussion

Large-scale public repositories contain decades of gene-expression data generated across multiple technologies, yet these datasets are rarely analyzed together because of substantial platform-specific differences. This disconnection limits statistical power, reduces reproducibility, and prevents biological discoveries that could emerge from integrative analyses. PXN addresses this challenge by providing a probabilistic framework that learns a shared representation of biological signal across technologies, enabling reliable integration of data from different platforms.

Across simulations and real datasets, PXN consistently produced more accurate cross-platform predictions than existing approaches. In multi-platform settings, its integrated training strategy improved normalization accuracy beyond that of individually trained models and enabled harmonization even between platforms with no direct overlap. Notably, our real-data analyses demonstrated that PXN can successfully harmonize even the most different technologies such as microarray and RNA-seq and enable them to be analyzed jointly without sacrificing statistical rigor. Differential-expression analyses based on PXN-normalized data retained high power and proper type I error control, outperforming both separate platform-specific meta-analysis (pcomb) and unnormalized integration. By enabling the joint analysis of legacy microarray cohorts and modern RNA-seq studies within a coherent statistical framework, PXN substantially expands the scope of biologically meaningful and statistically robust integrative genomics.

A major strength of PXN is its practical utility. Operating on a “learn once, apply everywhere” architecture, a single global model seamlessly translates data across any combination of supported technologies without sequential retraining. PXN is also computationally efficient: by exploiting Kronecker product structure in the covariance of multi-platform data, PXN reduces operations on extremely large matrices with dimension (*mK*) × (*mK*) to repeated operations of *m* × *m* matrices, which are further simplified using the Woodbury identity into low-dimensional updates involving *L* × *L* and *L* × *m* matrices (see Sections S1 – S3, *Supplementary Information*). This matrix-analytic design ensures numerical stability and allows PXN to handle tens of thousands of genes across many platforms, making it highly suitable for large-scale resources such as GEO, ArrayExpress, and TCGA.

PXN also has limitations that motivate several directions for future development. It assumes that the shared biological variation can be captured by a linear latent-factor model, which is effective in many transcriptomic settings but may not fully represent real-world gene–gene relationships. Extensions based on kernel PCA or deep encoder–decoder architectures could allow PXN to model more complex latent structure, though such nonlinear approaches would introduce challenges for interpretability that must be considered. Additionally, the current PXN probabilistic model is based on multivariate normality targeted for log-transformed bulk expression data. Adapting PXN to non-Gaussian models would make it applicable for other modalities such as single-cell RNA-seq, spatial assays, or other count-based data. These extensions represent promising avenues for enhancing PXN’s flexibility while preserving its core strengths in scalability and cross-platform harmonization.

In summary, PXN offers a robust, scalable, and biologically grounded solution for cross-technology harmonization. It allows researchers to combine data from vastly different platforms and realize the full potential of public gene-expression repositories.

## Methods

### Statistical Model

In this study, we consider a set of *N* biological samples sequenced on *K* technical platforms to produce expression profiles for *m* genes. The results are represented by gene expression matrices *Y*_*k*_ ∈ ℝ^*m*×*N*^, for *k* ∈ {1,…,*K*}. PXN models platform effects as gene-specific affine transformations applied to a shared underlying biological signal *Y*^(0)^:

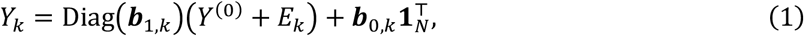

where ***b***_0, *k*_, ***b***_1,*k*_ ∈ ℝ^*m*^ are gene- and platform-specific intercepts and slopes; *E*_*k*_ ∈ ℝ^*m*×*N*^denotes independent measurement error with heteroskedastic (gene-specific) variance var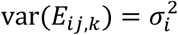. The underlying biological expression matrix *Y*^(0)^ ∈ ℝ^*m*×*N*^ is modeled by an extended PPCA framework:

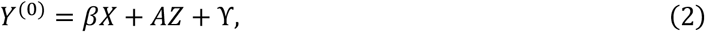

where *X* ∈ ℝ^*p*×*N*^ contains *p* covariates, β ∈ ℝ^*m*×*p*^ are linear coefficients that encode covariate effects. *Z* ∈ ℝ^*L*×N^ (*L* ≪ *m*) are latent factors capturing shared variation across genes, and *A* ∈ ℝ^*m*×*L*^ consists of gene-specific loadings. ϒ ∈ ℝ^m×N^ represents gene-specific variations not explained by *X* or *Z*, with heteroskedastic variances 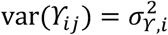.

This generative model provides a principled foundation for information-sharing across genes, covariate adjustment, and cross-platform prediction through conditional expectation. Identifiability constraints, variance normalization, and mean-centering conditions are described in *Supplementary Information* Section S1.

### Parameter Estimation

PXN parameters are estimated through a hybrid algorithm which combines closed-form updates with an EM-style blockwise optimization. Initial estimates of β, *A*, noise variances, and platform-specific affine parameters (***b***_0,k_, ***b***_1,k_) are obtained from regression- and conditional maximum likelihood estimators (*Supplementary Information* Section S2). These *closed-form* estimators provide stable and computationally efficient initialization that is essential for optimization in high-dimensional parameter spaces. We also implemented an optional refinement step using gradient descent to further improve likelihood-based parameter estimates. Users may choose between:

- **PXN (full)** — includes gradient-descent refinement for better accuracy
- **PXN-noGD** — closed-form estimation only; slightly less accurate but substantially faster

All large-matrix inversions are replaced by low-dimensional operations by exploiting the Kronecker structure in platform-level covariance matrices, which allows us to reduce operations on (*mK*) × (*mK*) matrices to repeated operations of *m* × *m* matrices. We further reduce these operations to computations of *L* × *m* and *L* × *L* matrices using the latent factor structure and the Woodbury identity. These matrix analysis techniques significantly improve numerical stability and scalability. Detailed derivations are provided in *Supplementary Information* Sections S2 – S3.

### Model-Based Cross-Platform Normalization

A key advantage of PXN is its ability to jointly leverage platform-specific parameters, clinical covariates, gene-specific variability, and latent factors shared across genes. After fitting the PXN model, we can use the following conditional expectation to normalize gene expression measurements from any source platform *k* to any target platform *k*′ without retraining:

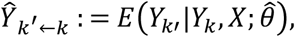

where 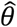 denotes the estimated model parameters. Note that 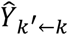is the best linear unbiased predictor (EBLUP) under the model assumptions which enjoys certain theoretical optimality properties (Henderson 1975, Robinson 1991). It also has a closed-form expression that allows efficient computation. An interesting point is that this estimator can be explicitly decomposed as 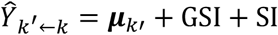, where 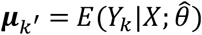 is the platform-specific mean; GSI (gene-specific information) depends only on quantities related to the *i*th genes; SI (shared information) captures multivariate correlation among genes through latent factors. This decomposition shows how PXN borrows information across genes via latent factors to achieve more accurate cross-platform normalization. Complete mathematical formulations and implementation details are provided in *Supplementary Information* Section S3.

### Multi-Platform Model Bridging

In many practical scenarios, the collection of relevant transcriptomic data often consists of several independent or partially overlapping platform pairs. For example, in **Figure 1**, the data collection can be divided into three *pairing groups*: AB, BC, and EF. Such structure is especially common in multi-center studies and in analyses that combine heterogeneous public datasets to increase statistical power. To use these data effectively, PXN provides a model bridging method that combines information across all available pairing groups. The result is a unified model which can consistently harmonize expression measures even between platforms that are not directly connected through shared samples (i.e., zero-shot normalization).

This model integration procedure starts with separate parameter estimations within each pairing group, yielding estimates that we collectively denote by 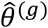for *g* = 1, … *G*. We then combine these estimates by using the following algorithm.

1. **Shared parameters** *β,Σ*_E_, and *Σ*_ϒ_. For these parameters shared across all platforms, we compute sample-size–weighted averages as the corresponding parameters for the integrated model.
2. **Latent factor covariance** *AA*′. We designed an integrative estimation procedure to estimate a common latent factor subspace using all available samples after properly removing platform-specific effects. This algorithm is computationally scalable and the resulting latent factor covariance matrix is guaranteed to be positive semi-definite.
3. **Platform-specific parameters *b***_0,*k*_ and ***b***_1,*k*_. For platform-specific parameters ***b***_0,*k*_ and ***b***_1,*k*_, we take the weighted average only across the subset of groups in which platform *k* appears.
4. **Estimation of the shared latent structure**. Compute the standardized residuals from all groups using the integrated parameter estimates computed in Step 2. Apply singular-value decomposition (SVD) to them to estimate a unified low-rank covariance estimate *ÂÂ*′. This step aligns the latent factors across groups and ensures a consistent biological representation across all platforms.

Complete mathematical derivations and implementation details are provided in *Supplementary Information* Section S4.

### Statistical Analysis

The primary quantitative metric for normalization accuracy was the Mean Squared Prediction Error (MSPE) between the projected and observed target expression values using 10-fold cross-validation. Downstream differential gene expression analyses (DGEA) were performed using Ordinary Least Squares (OLS) regression and Linear Mixed-Effects Models (LMER). Performance of DGEA as applied to various normalized data was evaluated using the True Positive Rate (Power), False Positive Rate (FPR), and the Area Under the Receiver Operating Characteristic Curve (AUC), averaged across 100 independent training-test data splits.

## Data Availability

All real datasets analyzed in this study are publicly available. The CellMiner transcriptomic profiles used in Real-data Demonstrations I and II were obtained from the NCI CellMiner resource, including multi-platform Affymetrix, Agilent, and RNA-seq expression datasets. Specifically, the processed microarray (Affymetrix and Agilent) and RNA-seq expression data were downloaded directly from the official CellMiner website. The TCGA dataset used in real-data demonstration III was downloaded from the TCGA Data Portal (Cancer Genome Atlas Research, Weinstein et al. 2013) and includes matched tumor/normal mRNA expression profiles from 583 breast cancer patients, quantified using both a microarray platform (Agilent G4502A) and an RNA-seq platform (GPL11154). All datasets were preprocessed using within-platform normalization and log2 transformation. Data preparation details are described in *Supplementary Information* Sections S5-S6.

To facilitate reproducibility, the processed CellMiner expression matrices and TCGA processed analysis inputs used in the analyses are provided with the PXN R package at https://github.com/zhiningsui/PXN. The platform and sample harmonization files, cross-validation splits, and summary results reported in

Tables 5–6 and Supplementary Tables S3 and S4 are available from the corresponding author upon reasonable request.

## Code Availability

The proposed method is implemented in the R package PXN, which is freely available at https://github.com/zhiningsui/PXN. The repository includes source code for model fitting, cross-platform prediction, model bridging, and processed analysis inputs.

## Ethics declarations

This study used only publicly available, de-identified data and did not require additional ethical approval.

## Author contributions

J.Z. and X.Q. conceived the experiments, Z.S. and D.Y. implemented the algorithms. Z.S., D.Y., and A.E. carried out simulations and real data analyses. All authors contributed to the writing of the manuscript and approved the final version for submission.

## References

Almudevar, A., L. B. Klebanov, X. Qiu, P. Salzman and A. Y. Yakovlev (2006). “Utility of correlation measures in analysis of gene expression.” NeuroRx 3(3): 384–395.

Alter, O., P. O. Brown and D. Botstein (2000). “Singular value decomposition for genome-wide expression data processing and modeling.” Proc Natl Acad Sci U S A 97(18): 10101–10106.

Benito, M., J. Parker, Q. Du, J. Wu, D. Xiang, C. M. Perou and J. S. Marron (2004). “Adjustment of systematic microarray data biases.” Bioinformatics 20(1): 105–114.

Borisov, N., I. Shabalina, V. Tkachev, M. Sorokin, A. Garazha, A. Pulin, Eremin, II and A. Buzdin (2019). “Shambhala: a platform-agnostic data harmonizer for gene expression data.” BMC Bioinformatics 20(1): 66.

Borisov, N., M. Sorokin, M. Zolotovskaya, C. Borisov and A. Buzdin (2022). “Shambhala-2: A Protocol for Uniformly Shaped Harmonization of Gene Expression Profiles of Various Formats.” Curr Protoc 2(5): e444.

Brazma, A., H. Parkinson, U. Sarkans, M. Shojatalab, J. Vilo, N. Abeygunawardena, E. Holloway, M. Kapushesky, P. Kemmeren, G. G. Lara, A. Oezcimen, P. Rocca-Serra and S. A. Sansone (2003). “ArrayExpress--a public repository for microarray gene expression data at the EBI.” Nucleic Acids Res 31(1): 68–71.

Cancer Genome Atlas Research, N., J. N. Weinstein, E. A. Collisson, G. B. Mills, K. R. Shaw, B. A. Ozenberger, K. Ellrott, I. Shmulevich, C. Sander and J. M. Stuart (2013). “The Cancer Genome Atlas Pan-Cancer analysis project.” Nat Genet 45(10): 1113–1120.

Consortium, M. (2006). “The MicroArray Quality Control (MAQC) project shows inter- and intraplatform reproducibility of gene expression measurements.” Nature Biotechnology 24(9): 1151–1161.

Consortium, S. M.-I. (2014). “A comprehensive assessment of RNA-seq accuracy, reproducibility and information content by the Sequencing Quality Control Consortium.” Nature Biotechnology 32(9): 903–914.

Cui, H., C. Wang, H. Maan, K. Pang, F. Luo, N. Duan and B. Wang (2024). “scGPT: toward building a foundation model for single-cell multi-omics using generative AI.” Nat Methods 21(8): 1470–1480.

Cui, Z., Y. Liu, J. Zhang and X. Qiu (2021). “Super-delta2: an enhanced differential expression analysis procedure for multi-group comparisons of RNA-seq data.” Bioinformatics 37(17): 2627–2636.

de la Fuente, A. (2010). “From ‘differential expression’ to ‘differential networking’ - identification of dysfunctional regulatory networks in diseases.” Trends Genet 26(7): 326–333.

Edgar, R., M. Domrachev and A. E. Lash (2002). “Gene Expression Omnibus: NCBI gene expression and hybridization array data repository.” Nucleic Acids Res 30(1): 207–210.

Efron, B. (2007). “Correlation and large-scale simultaneous significance testing.” Journal of the American Statistical Association 102(477): 93–103.

Henderson, C. R. (1975). “Best linear unbiased estimation and prediction under a selection model.” Biometrics 31(2): 423–447.

Hong, F. and R. Breitling (2008). “A comparison of meta-analysis methods for detecting differentially expressed genes in microarray experiments.” Bioinformatics 24(3): 374–382.

Ideker, T., V. Thorsson, J. A. Ranish, R. Christmas, J. Buhler, J. K. Eng, R. Bumgarner, D. R. Goodlett, R. Aebersold and L. Hood (2001). “Integrated genomic and proteomic analyses of a systematically perturbed metabolic network.” Science 292(5518): 929–934.

Irizarry, R. A., D. Warren, F. Spencer, I. F. Kim, S. Biswal, B. C. Frank, E. Gabrielson, J. G. Garcia, J. Geoghegan, G. Germino, C. Griffin, S. C. Hilmer, E. Hoffman, A. E. Jedlicka, E. Kawasaki, F. Martinez-Murillo, L. Morsberger, H. Lee, D. Petersen, J. Quackenbush, A. Scott, M. Wilson, Y. Yang, S. Q. Ye and W. Yu (2005). “Multiple-laboratory comparison of microarray platforms.” Nat Methods 2(5): 345–350.

Johnson, W. E., C. Li and A. Rabinovic (2007). “Adjusting batch effects in microarray expression data using empirical Bayes methods.” Biostatistics 8(1): 118–127.

Kedzierska, K. Z., L. Crawford, A. P. Amini and A. X. Lu (2023). “Assessing the limits of zero-shot foundation models in single-cell biology.” BioRxiv: 2023.2010.2016.561085.

Leek, J. T. and J. D. Storey (2007). “Capturing heterogeneity in gene expression studies by surrogate variable analysis.” PLoS Genet 3(9): 1724–1735.

Li, K. C. (2002). “Genome-wide coexpression dynamics: theory and application.” Proc Natl Acad Sci U S A 99(26): 16875–16880.

Liu, Y., J. Zhang and X. Qiu (2017). “Super-delta: a new differential gene expression analysis procedure with robust data normalization.” BMC Bioinformatics 18(1): 582.

Moretto, M., P. Sonego, A. B. Villasenor-Altamirano and K. Engelen (2019). “First step toward gene expression data integration: transcriptomic data acquisition with COMMAND>_.” BMC Bioinformatics 20(1): 54.

Nagalakshmi, U., Z. Wang, K. Waern, C. Shou, D. Raha, M. Gerstein and M. Snyder (2008). “The transcriptional landscape of the yeast genome defined by RNA sequencing.” Science 320(5881): 1344–1349.

Qiu, X., A. I. Brooks, L. Klebanov and N. Yakovlev (2005). “The effects of normalization on the correlation structure of microarray data.” BMC Bioinformatics 6: 120.

Qiu, X., R. Hu and Z. Wu (2014). “Evaluation of bias-variance trade-off for commonly used post-summarizing normalization procedures in large-scale gene expression studies.” PLoS One 9(6): e99380.

Qiu, X., L. Klebanov and A. Yakovlev (2005). “Correlation between gene expression levels and limitations of the empirical bayes methodology for finding differentially expressed genes.” Stat Appl Genet Mol Biol 4: Article 34.

Qiu, X., H. Wu and R. Hu (2013). “The impact of quantile and rank normalization procedures on the testing power of gene differential expression analysis.” BMC Bioinformatics 14: 124.

Robinson, G. K. (1991). “That BLUP is a good thing: the estimation of random effects.” Statistical science: 15–32.

Schena, M., D. Shalon, R. W. Davis and P. O. Brown (1995). “Quantitative monitoring of gene expression patterns with a complementary DNA microarray.” Science 270(5235): 467–470.

Sielemann, K., A. Hafner and B. Pucker (2020). “The reuse of public datasets in the life sciences: potential risks and rewards.” PeerJ 8: e9954.

Tipping, M. E. and C. M. Bishop (1999). “Probabilistic principal component analysis.” Journal of the Royal Statistical Society Series B: Statistical Methodology 61(3): 611–622.

Tseng, G. C., D. Ghosh and E. Feingold (2012). “Comprehensive literature review and statistical considerations for microarray meta-analysis.” Nucleic Acids Res 40(9): 3785–3799.

Wang, Z., M. Gerstein and M. Snyder (2009). “RNA-Seq: a revolutionary tool for transcriptomics.” Nat Rev Genet 10(1): 57–63.

Zhang, S., J. Shao, D. Yu, X. Qiu and J. Zhang (2020). “MatchMixeR: a cross-platform normalization method for gene expression data integration.” Bioinformatics 36(8): 2486–2491.

Zhao, S., W. P. Fung-Leung, A. Bittner, K. Ngo and X. Liu (2014). “Comparison of RNA-Seq and microarray in transcriptome profiling of activated T cells.” PLoS One 9(1): e78644.

