## Supplementary Information for "PXN Unlocks the Power of Public Gene Expression Data Through Cross-Technology Integration"

#### S1 PXN Model

In this study, we denote by $Y_{ij,k}$ the observed expression level of gene $i = 1, \ldots, m$ in paired sample $j = 1, \ldots, N$, measured on platform $k = 1, \ldots, K$. For each platform $k$, the data form a matrix $Y_k \in \mathbb{R}^{m \times N}$, with rows corresponding to genes and columns to samples:

$$\begin{pmatrix} Y_{11,1} & Y_{12,1} & \cdots & Y_{1N,1} \\ Y_{21,1} & Y_{22,1} & \cdots & Y_{2N,1} \\ \vdots & \vdots & \ddots & \vdots \\ Y_{m1,1} & Y_{m2,1} & \cdots & Y_{mN,1} \end{pmatrix}, \ldots, \begin{pmatrix} Y_{11,K} & Y_{12,K} & \cdots & Y_{1N,K} \\ Y_{21,K} & Y_{22,K} & \cdots & Y_{2N,K} \\ \vdots & \vdots & \ddots & \vdots \\ Y_{m1,K} & Y_{m2,K} & \cdots & Y_{mN,K} \end{pmatrix}$$

If the samples are perfectly matched across $K$ platforms, we may vertically stack these platform-specific expression matrices to form a single aggregated matrix $Y$:

$$Y := \left(\begin{array}{cccc} Y_{11,1} & Y_{12,1} & \cdots & Y_{1N,1} \\ Y_{21,1} & Y_{22,1} & \cdots & Y_{2N,1} \\ \vdots & \vdots & \ddots & \vdots \\ Y_{m1,1} & Y_{m2,1} & \cdots & Y_{mN,1} \\ \hline Y_{11,2} & Y_{12,2} & \cdots & Y_{1N,2} \\ Y_{21,2} & Y_{22,2} & \cdots & Y_{2N,2} \\ \vdots & \vdots & \ddots & \vdots \\ Y_{m1,2} & Y_{m2,2} & \cdots & Y_{mN,2} \\ \hline \vdots & \vdots & \vdots & \vdots \\ \vdots & \vdots & \vdots & \vdots \\ \hline Y_{11,K} & Y_{12,K} & \cdots & Y_{1N,K} \\ Y_{21,K} & Y_{22,K} & \cdots & Y_{2N,K} \\ \vdots & \vdots & \ddots & \vdots \\ Y_{m1,K} & Y_{m2,K} & \cdots & Y_{mN,K} \end{array}\right) \in \mathbb{R}^{mK\times N}.$$

We define a set of *latent standardized* gene expressions, denoted as $Y_{ij}^{(0)}$, representing the true underlying expression level of gene $i$ in sample $j$, free from platform effects and measurement error. These values are assumed to be driven by $L$ unobserved latent factors. The term "standardized" refers to the constraint that each gene's expression vector across samples has unit variance—that is, each row of the matrix $Y^{(0)} \in \mathbb{R}^{m\times N}$ has standard deviation equal to 1. This standardization is imposed for identifiability (see Section S1.B.1) and does not imply the true biological variability for every gene equals one.

The platform-independent true gene expression $Y^{(0)}$ captures the underlying biological signal shared across all platforms. On each platform, the observed gene expressions are affine (linear plus the intercept) transformations of $Y_{ij}^{(0)}$, plus measurement error.

#### S1.A Model for Latent Standardized Gene Expressions

We assume that each sample is associated with a set of covariates, such as Cell-line, Sex, Age, etc, denoted as $\mathbf{X}_j = (1, X_{2j}, \ldots, X_{pj})' \in \mathbb{R}^p$. In addition, let $\mathbf{Z}_j = (Z_{1j}, \cdots, Z_{Lj})' \in \mathbb{R}^L$ represent the $L$ unobservable latent factors *shared across all genes* for sample $j$. Let $X = (\mathbf{X}_1 \ \cdots \ \mathbf{X}_N) \in \mathbb{R}^{p\times N}$ and $Z = (\mathbf{Z}_1 \ \cdots \ \mathbf{Z}_N) \in \mathbb{R}^{L\times N}$ denote the matrix of covariates and the matrix of latent factors, respectively, across $N$ samples. These latent factors are assumed to be independent standard normal:

$$\mathbf{Z}_j \sim N(\mathbf{0}_L, I_L), \quad j = 1, ..., N, \tag{S1.1}$$

which implies that the column-stacked vector $\text{vec}(Z) \in \mathbb{R}^{LN}$ satisfies:

$$\text{vec}(Z) \sim N(0_{LN}, I_{LN}).$$

We assume that the covariates (observable) and the latent factors (unobservable) "drive" the variations in *latent standardized gene expressions*. So, we model the latent standardized gene expression vector for sample $j$ as

$$\mathbf{Y}_j^{(0)} = \beta \mathbf{X}_j + A\mathbf{Z}_j + \boldsymbol{\upsilon}_j, \tag{S1.2}$$

where $\mathbf{Y}_j^{(0)} = (Y_{1j}^{(0)}, \ldots, Y_{mj}^{(0)})' \in \mathbb{R}^m$ is the $j$-th column of $Y^{(0)}$, $\beta \in \mathbb{R}^{m\times p}$ links $\mathbf{X}_j$ to $\mathbf{Y}_j^{(0)}$, $A \in \mathbb{R}^{m\times L}$ is the factor loading matrix linking latent factors to the shared variation in $\mathbf{Y}_j^{(0)}$, and $\boldsymbol{\upsilon}_j \in \mathbb{R}^m$ represents gene-specific residual variation not explained by the latent structure. We assume independent residual noise across genes and samples:

$$\boldsymbol{\upsilon}_j \sim N(\mathbf{0}_m, \Sigma_\Upsilon), \quad \text{with} \quad \Sigma_\Upsilon = \text{Diag}(\sigma^2_{\Upsilon,1}, \ldots, \sigma^2_{\Upsilon,m}). \tag{S1.3}$$

Therefore, based on Equation (S1.2), we have

$$\mathbf{Y}_j^{(0)} \sim N(\beta \mathbf{X}_j, \Sigma^{(0)}), \quad \text{where} \quad \Sigma^{(0)} := AA' + \Sigma_\Upsilon. \tag{S1.4}$$

Stacking all $N$ samples and defining $\Upsilon = (\boldsymbol{\upsilon}_1 \ \cdots \ \boldsymbol{\upsilon}_N) \in \mathbb{R}^{m\times N}$ and $Y^{(0)} = \begin{pmatrix}\mathbf{Y}_1^{(0)} & \cdots & \mathbf{Y}_N^{(0)}\end{pmatrix}$, we obtain the full latent model:

$$\begin{gathered} Y^{(0)} = \beta X + AZ + \Upsilon, \\ \text{vec}(Z) \sim N(\mathbf{0}_{LN}, I_{LN}), \quad \text{vec}(\Upsilon) \sim N(\mathbf{0}_{mN}, I_N \otimes \Sigma_\Upsilon). \\ \text{vec}(Y^{(0)}) \sim N((I_N \otimes \beta)\text{vec}(X),\ I_N \otimes \Sigma^{(0)}). \end{gathered} \tag{S1.5}$$

The latent variable $Y^{(0)}$ serves as a conceptual tool to help us derive the covariance between the observed gene expressions ($Y_k$), which leads to the conditional expectations between them that are needed for cross-platform normalization.

##### S1.A.1 Distribution of Latent Standardized Gene Expressions

To clarify the distribution of the latent standardized gene expressions $Y^{(0)}$, we recall that while $Z \in \mathbb{R}^{L\times N}$ is matrix-valued, distributions are defined over vectorized forms. Thus, we work with:

$$\text{vec}(Z) \sim N(\mathbf{0}_{LN}, I_{LN}).$$

Focusing on the random component of $Y^{(0)} = AZ$, we have:

$$\text{vec}(Y^{(0)}) = \text{vec}(AZ) = (I_N \otimes A)\text{vec}(Z), \tag{S1.6}$$

and hence,

$$\text{Var}[\text{vec}(Y^{(0)})] = \text{Var}[(I_N \otimes A)\text{vec}(Z)] = (I_N \otimes A)(I_{LN})(I_N \otimes A)' = I_N \otimes (AA'). \tag{S1.7}$$

Adding the residual noise $\Upsilon$, which is independent of $Z$, the total variance becomes:

$$\text{Var}[\text{vec}(Y^{(0)})] = I_N \otimes (AA' + \Sigma_\Upsilon) = I_N \otimes \Sigma^{(0)}. \tag{S1.8}$$

Therefore, the vectorized latent standardized gene expressions follow a normal distribution:

$$\text{vec}(Y^{(0)}) \sim N((I_N \otimes \beta)\text{vec}(X),\ I_N \otimes \Sigma^{(0)}). \tag{S1.9}$$

#### S1.B Model for Observed Gene Expressions

The observed expression on platform $k$ is modeled as an affine transformation of the true latent expression, with added measurement error:

$$\begin{aligned}
Y_{ij,k} &= b_{1i,k}\left(Y_{ij}^{(0)} + E_{ij,k}\right) + b_{0i,k} \\
&= b_{1i,k}\left(\beta_{i1} + \sum_{r=2}^{p}\beta_{ir}X_{rj} + \sum_{l=1}^{L}a_{il}Z_{lj} + \Upsilon_{ij} + E_{ij,k}\right) + b_{0i,k} \\
&= b_{1i,k}\beta_{i1} + b_{0i,k} + \sum_{r=2}^{p}b_{1i,k}\beta_{ir}X_{rj} + \sum_{l=1}^{L}b_{1i,k}a_{il}Z_{lj} + b_{1i,k}\Upsilon_{ij} + b_{1i,k}E_{ij,k} \\
&= \sum_{r=1}^{p}c_{ir,k}X_{rj} + \sum_{l=1}^{L}d_{il,k}Z_{lj} + b_{1i,k}\Upsilon_{ij} + b_{1i,k}E_{ij,k}, \\
c_{ir,k} &:= \begin{cases} b_{1i,k}\beta_{i1} + b_{0i,k}, & r = 1, \\ b_{1i,k}\beta_{ir}, & r = 2, \ldots, p, \end{cases} \qquad d_{il,k} := b_{1i,k}a_{il}.
\end{aligned} \tag{S1.10}$$

where $b_{0i,k}$ and $b_{1i,k}$ are platform-specific intercepts and slopes for gene $i$, $E_{ij,k}$ represents intrinsic measurement error. The resulting term $b_{1i,k}E_{ij,k}$ accounts for extrinsic error in the observed value $Y_{ij,k}$.

In theory, the variance of $E_{ij,k}$ may depend on the genes and platforms. However, such model includes too many unknown variance parameters ($mK$ of them) for $\text{var}(E_{ij,k})$), which makes the subsequent parameter estimation difficult. On the other hand, it is not realistic to assume that all genes have exactly the same level of measurement error. As a compromise, we impose the following *Proportional Variance Assumption*. We assume that for a gene $i$, $E_{ij,k}$ are independent with gene-specific variances shared across all technical platforms: $\text{Var}(E_{ij,k}) = \sigma_i^2$. This results in a diagonal covariance matrix for the intrinsic errors, $\Sigma_E = \text{Diag}(\sigma_i^2) \in \mathbb{R}^{m\times m}$. Consequently, the variance of the extrinsic measurement error is proportional to $b_{1i,k}$.

It is convenient to express the model at the platform level. Let $Y_k \in \mathbb{R}^{m\times N}$ denote the observed gene expression matrix for platform $k$, with columns indexed by samples. Then, the matrix version of Equation (S1.10) becomes:

$$\begin{aligned}
Y_k &= \text{Diag}(\mathbf{b}_{1,k})\left(Y^{(0)} + E_k\right) + \mathbf{b}_{0,k}\mathbf{1}_N' \\
&= \text{Diag}(\mathbf{b}_{1,k})\left(\beta X + AZ + \Upsilon + E_k\right) + \mathbf{b}_{0,k}\mathbf{1}_N \\
&= C_kX + \text{Diag}(\mathbf{b}_{1,k})\left(AZ + \Upsilon + E_k\right), \\
C_k &= \text{Diag}(\mathbf{b}_{1,k})\beta + \left(\mathbf{b}_{0,k}\,\middle|\,\mathbf{0}_{m\times(p-1)}\right), \quad \text{Diag}(\mathbf{b}_{1,k}) := \begin{pmatrix} b_{11,k} & & \\ & \ddots & \\ & & b_{1m,k} \end{pmatrix},
\end{aligned} \tag{S1.11}$$

where $\mathbf{b}_{0,k} = (b_{01,k}, \ldots, b_{0m,k})' \in \mathbb{R}^m$ and $\mathbf{b}_{1,k} = (b_{11,k}, \ldots, b_{1m,k})' \in \mathbb{R}^m$ denote vectors of intercepts and slopes for all genes specific to platform $k$. Here, $Z$, $\Upsilon$, and $E_k$ are assumed to be independent of each other, and intrinsic measurement errors ($E_{ij,k}$) are all independent but not necessarily identically distributed.

The decomposition of $Y_k$ in Equation (S1.11) includes a mean component $C_k X$, shared latent variation $\text{Diag}(\mathbf{b}_{1,k})AZ$, gene-specific variation $\text{Diag}(\mathbf{b}_{1,k})\Upsilon$, and measurement error $\text{Diag}(\mathbf{b}_{1,k})E_k$. This model can be considered as a combination of standard regression model (which only has the first two terms) and the probabilistic PCA model (which has the shared and gene-specific variations).

##### S1.B.1 Solving the Identifiability Issues

Because we can not directly observe the true gene expressions ($Y^{(0)}$), not all parameters in Equation (S1.11) are *identifiable*. Thus, identifiability constraints are imposed on $Y^{(0)}$ to ensure that not all parameters are free to vary.

Let $Y_k^{(0)} := \text{Diag}(\mathbf{b}_{1,k})Y^{(0)} + \mathbf{b}_{0,k}\mathbf{1}'_N$. It represents the noise-free gene expression measured on the $k$th platform and is an affine transformation of $Y^{(0)}$ by assumption. Let $\tilde{Y}^{(0)} := \text{Diag}(\mathbf{h}_1)Y^{(0)} + \mathbf{h}_0\mathbf{1}'_N$ be an alternative choice of the true gene expression, where $\mathbf{h}_0, \mathbf{h}_1$ are two arbitrary vectors in $\mathbb{R}^m$. If we choose $\tilde{\mathbf{b}}_{0,k} := \mathbf{b}_{0,k} - \mathbf{b}_{1,k} \circ \mathbf{h}_1^{-1} \circ \mathbf{h}_0$, $\tilde{\mathbf{b}}_{1,k} := \mathbf{b}_{1,k} \circ \mathbf{h}_1^{-1}$, and $\tilde{E}_k := \text{Diag}(\mathbf{h}_1)E_k$ (here "$\circ$" stands for the Hadamard product), we have

$$\text{Diag}(\tilde{\mathbf{b}}_{1,k}) \left(\tilde{Y}^{(0)} + \tilde{E}_k\right) + \tilde{\mathbf{b}}_{0,k}\mathbf{1}'_N = \text{Diag}(\mathbf{b}_{1,k}) \left(Y^{(0)} + E_k\right) + \mathbf{b}_{0,k}\mathbf{1}'_N = Y_k. \tag{S1.12}$$

In other words, this alternative representation is mathematically equivalent to the original representation of the observed gene expressions $Y_k$.

One way to resolve this problem is to impose mathematical constraints to the expectation and covariance of $Y^{(0)}$. In the proposed algorithm, we will use the *standardized* gene expressions as a means to obtain an initial set of parameter estimators. Specifically, for each gene, we impose the following constraints:

$$\text{var}\left(Y_{ij}^{(0)} + E_k\right) = 1 \Longrightarrow |A_{i\cdot}|^2 + \sigma_{\Upsilon,i}^2 + \sigma_i^2 = 1. \tag{S1.13a}$$

$$E\overline{\tilde{Y}_i^{(0)}} = 0 \Longrightarrow \beta\bar{X} = 0_m. \tag{S1.13b}$$

Here $\bar{\cdot}$ represents average in the direction of subjects: $\overline{\tilde{Y}_i^{(0)}} := \frac{1}{N}\sum_{j=1}^N \tilde{Y}_{ij}^{(0)} = \frac{1}{N}\tilde{Y}_{ij}^{(0)}\mathbf{1}_N$, $\bar{X} := \frac{1}{N}\sum_{j=1}^N X_{pj} = \frac{1}{N}X\mathbf{1}_N$. Constraint (S1.13b) can also be written as: $\beta_{i1} + \sum_{r=2}^p \beta_{ir}\bar{X}_r = 0$ for all $i = 1, 2, \ldots, m$.

#### S1.C The Joint Log-Likelihood Function

To estimate the parameters in the Model (S1.11), we adopt a likelihood-based approach. We derive the joint log-likelihood function for the observed gene expression data $Y_k$ across all platforms $k = 1, ..., K$. The goal is to estimate parameters $\theta = \{\mathbf{b}_{0,k}, \mathbf{b}_{1,k}, \sigma^2, \sigma_\Upsilon^2, A\}$.

##### S1.C.1 Sample-wise Expression Vector

Let $\mathbf{Y}_j \in \mathbb{R}^{mK}$ denote the vertically stacked vector of observed expressions for sample $j$ across all platforms:

$$\mathbf{Y}_j := \begin{pmatrix} \mathbf{Y}_{j,1} \\ \hline \mathbf{Y}_{j,2} \\ \hline \vdots \\ \hline \mathbf{Y}_{j,K} \end{pmatrix} \in \mathbb{R}^{mK}, \quad \text{where} \quad \mathbf{Y}_{j,k} \in \mathbb{R}^m.$$

Define similarly stacked slope and intercept vectors: $\mathbf{b}_0 = (\mathbf{b}'_{0,1} \ \dots \ \mathbf{b}'_{0,K})' \in \mathbb{R}^{mK}$ and $\mathbf{b}_1 = (\mathbf{b}'_{1,1} \ \dots \ \mathbf{b}'_{1,K})' \in \mathbb{R}^{mK}$. Let $B_1 := \mathrm{Diag}(\mathbf{b}_1) \in \mathbb{R}^{mK \times mK}$ denote the block-diagonal matrix representing the platform-specific slopes:

$$B_1 = \begin{pmatrix} \mathrm{Diag}(\mathbf{b}_{1,1}) & & & & \\ & \mathrm{Diag}(\mathbf{b}_{1,2}) & & & \\ & & \ddots & & \\ & & & \mathrm{Diag}(\mathbf{b}_{1,k}) & & \\ & & & & \ddots & \\ & & & & & \mathrm{Diag}(\mathbf{b}_{1,K}) \end{pmatrix} \tag{S1.14}$$

##### S1.C.2 Distribution of $\mathbf{Y}_j$

Since the latent standardized expression $Y^{(0)} = \beta X + AZ + \Upsilon$ is shared across platforms, the expectation of $\mathbf{Y}_j$ is

$$\boldsymbol{\mu}_{Y_j} := E\mathbf{Y}_j = B_1(\mathbf{1}_K \otimes \beta \mathbf{X}_j) + \mathbf{b}_0 \tag{S1.15}$$

We assume that the covariance matrix of the latent expressions ($\Sigma^{(0)} = AA' + \Sigma_\Upsilon$) to be the same across all samples. Let $\Sigma_Y \in \mathbb{R}^{mK \times mK}$ denote the covariance matrix of $\mathbf{Y}_j$. Based on the Model (S1.11), $\Sigma_Y$ can be expressed as

$$\begin{aligned} \Sigma_Y &= B'_1 \left[ \begin{pmatrix} \Sigma^{(0)} & \dots & \Sigma^{(0)} \\ \vdots & \ddots & \vdots \\ \Sigma^{(0)} & \dots & \Sigma^{(0)} \end{pmatrix} + \begin{pmatrix} \Sigma_E & & \\ & \ddots & \\ & & \Sigma_E \end{pmatrix} \right] B_1 \\ &= B_1 \left[ J_K \otimes (AA' + \Sigma_\Upsilon) + I_K \otimes \Sigma_E \right] B_1. \end{aligned} \tag{S1.16}$$

Here $\Sigma_E \in \mathbb{R}^{m \times m}$ is a diagonal matrix such that $\Sigma_{E,ii} = \sigma_i^2$, the intrinsic variance of measurement errors for the $i$th gene.

Therefore, based on Equations (S1.15) and (S1.16), we have

$$\mathbf{Y}_j \sim N\left(B_1(\mathbf{1}_K \otimes \beta \mathbf{X}_j) + \mathbf{b}_0,\ B_1 \left[J_K \otimes (AA' + \Sigma_\Upsilon) + I_K \otimes \Sigma_E\right] B_1\right). \tag{S1.17}$$

To compute the inverse of $\Sigma_Y$, define

$$\tilde{A} := \Sigma_\Upsilon^{-1/2} A, \qquad \tilde{B}_1 := B_1 \left( I_K \otimes \Sigma_E^{1/2} \right), \qquad G := \mathbf{1}_K \otimes (\Sigma_E^{-1/2} \Sigma_\Upsilon^{1/2}).$$

Then, we have

$$\begin{aligned}\Sigma_Y &= \tilde{B}_1 \left[ J_K \otimes \left( \Sigma_E^{-1/2} \Sigma_\Upsilon^{1/2} (\tilde{A}\tilde{A}' + I_m) \Sigma_\Upsilon^{1/2} \Sigma_E^{-1/2} \right) + I_{Km} \right] \tilde{B}_1 \\ &= \tilde{B}_1 \left[ G(\tilde{A}\tilde{A}' + I_m) G' + I_{Km} \right] \tilde{B}_1.\end{aligned} \tag{S1.18}$$

Let $\tilde{A} := UDV'$ be the thin SVD of $\tilde{A}$. Here $U \in \mathbb{R}^{m \times L}$ is a semi-orthogonal matrix, $D = \mathrm{Diag}(d_l) \in \mathbb{R}^{L \times L}$ is a diagonal matrix, and $V \in O(L)$. We have $\tilde{A}\tilde{A}' = UD^2U'$ and

$$\left| \tilde{A}\tilde{A}' + I_m \right| = \prod_{l=1}^{L} \left( 1 + d_l^2 \right), \qquad \left( \tilde{A}\tilde{A}' + I_m \right)^{-1} = I_m - U \left( D^{-2} + I_L \right)^{-1} U'.$$

$$\begin{aligned}\left| \left( \tilde{A}\tilde{A}' + I_m \right)^{-1} + G'G \right| &= \left| I_m + K\Sigma_E^{-1}\Sigma_\Upsilon - U \left( D^{-2} + I_L \right)^{-1} U' \right| \\ &= \left| D^{-2} + I_L - U'(I_m + K\Sigma_E^{-1}\Sigma_\Upsilon)^{-1} U \right| \cdot \frac{\left| I_m + K\Sigma_E^{-1}\Sigma_\Upsilon \right|}{|D^{-2} + I_L|} \\ &= C \prod_{i=1}^{m} \frac{\sigma_i^2 + K\sigma_{\Upsilon,i}^2}{\sigma_i^2}. \\ C &:= \frac{\left| D^{-2} + I_L - U'(I_m + K\Sigma_E^{-1}\Sigma_\Upsilon)^{-1} U \right|}{|D^{-2} + I_L|}.\end{aligned}$$

Note that in the above equation, $D^{-2} + I_L$ and $D^{-2} + I_L - U'(I_m + K\Sigma_E^{-1}\Sigma_\Upsilon)^{-1}U$ are both small ($L \times L$) matrices so $C$ is easy to compute numerically.

Next, we have

$$\begin{aligned}\left| G(\tilde{A}\tilde{A}' + I_m)G' + I_{Km} \right| &= \left| \left( \tilde{A}\tilde{A}' + I_m \right)^{-1} + G'G \right| \left| \tilde{A}\tilde{A}' + I_m \right| \\ &= C \cdot \prod_{i=1}^{m} \frac{\sigma_i^2 + K\sigma_{\Upsilon,i}^2}{\sigma_i^2} \cdot \prod_{l=1}^{L} (1 + d_l^2).\end{aligned}$$

$$\begin{aligned}|\Sigma_Y| &= C\, |B_1|^2 \, |\Sigma_E|^K \cdot \prod_{i=1}^{m} \frac{\sigma_i^2 + K\sigma_{\Upsilon,i}^2}{\sigma_i^2} \prod_{l=1}^{L} (1 + d_l^2) \\ &= C \left( \prod_{i,k} b_{1i,k}^2 \right) \left( \prod_{i=1}^{m} \sigma_i^2 \right)^{K-1} \cdot \prod_{i=1}^{m} (\sigma_i^2 + K\sigma_{\Upsilon,i}^2) \prod_{l=1}^{L} (1 + d_l^2).\end{aligned} \tag{S1.19}$$

$$\begin{aligned}\log |\Sigma_Y| &= \log C + 2 \sum_{i,k} \log b_{1i,k} + (K-1) \sum_{i=1}^{m} \log \sigma_i^2 \\ &\quad + \sum_{i=1}^{m} \log(\sigma_i^2 + K\sigma_{\Upsilon,i}^2) + \sum_{l=1}^{L} \log(1 + d_l^2).\end{aligned} \tag{S1.20}$$

More computations show

$$\begin{aligned}\left[G(\tilde{A}\tilde{A}'+I_m)G'+I_{Km}\right]^{-1} &= I_{km} - G\left(\left[\tilde{A}\tilde{A}'+I_m\right]^{-1}+G'G\right)^{-1}G' \\ &= I_{km} - G\left(I_m + K\Sigma_E^{-1}\Sigma_\Upsilon - U\left(D^{-2}+I_L\right)^{-1}U'\right)^{-1}G'.\end{aligned} \tag{S1.21}$$

Let $H := \left(I_m + K\Sigma_E^{-1}\Sigma_\Upsilon\right)^{-1} = \mathrm{Diag}\left(\frac{\sigma_i^2}{K\sigma_{\Upsilon,i}^2+\sigma_i^2}\right)$. We have

$$\begin{aligned} GH &= \left[\mathbf{1}_K \otimes (\Sigma_E^{-1/2}\Sigma_\Upsilon^{1/2})\right]\mathrm{Diag}\left(\frac{\sigma_i^2}{K\sigma_{\Upsilon,i}^2+\sigma_i^2}\right) = \mathbf{1}_K \otimes \mathrm{Diag}\left(\frac{\sigma_i\sigma_{\Upsilon,i}}{K\sigma_{\Upsilon,i}^2+\sigma_i^2}\right). \\ \tilde{B}_1^{-2} &= B_1^{-2}\left[I_K \otimes \Sigma_E^{-1}\right], \qquad \tilde{B}_1^{-1}GH = B_1^{-1}\left(\mathbf{1}_K \otimes \mathrm{Diag}\left(\frac{\sigma_{\Upsilon,i}}{K\sigma_{\Upsilon,i}^2+\sigma_i^2}\right)\right).\end{aligned} \tag{S1.22}$$

$$\begin{aligned} &(\tilde{B}_1^{-1}GH)\left(I_m + K\Sigma_E^{-1}\Sigma_\Upsilon\right)(\tilde{B}_1^{-1}GH)' \\ &= B_1^{-1}\left[J_K \otimes \mathrm{Diag}\left(\frac{\sigma_{\Upsilon,i}^2}{(K\sigma_{\Upsilon,i}^2+\sigma_i^2)^2}\cdot\left(1+\frac{K\sigma_{\Upsilon,i}^2}{\sigma_i^2}\right)\right)\right]B_1^{-1} \\ &= B_1^{-1}\left[J_K \otimes \mathrm{Diag}\left(\frac{\sigma_{\Upsilon,i}^2}{\sigma_i^2(K\sigma_{\Upsilon,i}^2+\sigma_i^2)}\right)\right]B_1^{-1}.\end{aligned}$$

$$\left(I_m + K\Sigma_E^{-1}\Sigma_\Upsilon - U\left(D^{-2}+I_L\right)^{-1}U'\right)^{-1} = H + HU\underbrace{\left(D^{-2}+I_L-U'HU\right)^{-1}}_{M}U'H. \tag{S1.23}$$

$$\begin{aligned}\left[G(\tilde{A}\tilde{A}'+I_m)G'+I_{Km}\right]^{-1} &= I_{km} - GH\left[H^{-1}+UMU'\right](GH)' \\ &= I_{km} - GH\left[I_m + K\Sigma_E^{-1}\Sigma_\Upsilon + UMU'\right](GH)'.\end{aligned}$$

Therefore

$$\begin{aligned}\Sigma_Y^{-1} &= \tilde{B}_1^{-1}\left[G(\tilde{A}\tilde{A}'+I_m)G'+I_{Km}\right]^{-1}\tilde{B}_1^{-1} \\ &= \tilde{B}_1^{-2} - \tilde{B}_1^{-1}GH\left[I_m + K\Sigma_E^{-1}\Sigma_\Upsilon + UMU'\right](\tilde{B}_1^{-1}GH)' \\ &= B_1^{-1}\left[I_K \otimes \Sigma_E^{-1} - J_K \otimes W\right]B_1^{-1}. \\ \tilde{B}_1^{-1} &= B_1^{-1}\left(I_K \otimes \Sigma_E^{-1/2}\right), \qquad \tilde{U} := \mathrm{Diag}\left(\frac{\sigma_{\Upsilon,i}}{K\sigma_{\Upsilon,i}^2+\sigma_i^2}\right)U. \\ W &:= \mathrm{Diag}\left(\frac{\sigma_{\Upsilon,i}^2}{\sigma_i^2(K\sigma_{\Upsilon,i}^2+\sigma_i^2)}\right) + \tilde{U}M\tilde{U}'.\end{aligned} \tag{S1.24}$$

Based on Equations (S1.17), (S1.19) and (S1.24), the grand log-likelihood function of all data is

$$\begin{aligned} -2\ell &= \underbrace{mNK \log 2\pi + N \log \det(\Sigma_Y)}_{\text{Term}_1} + \sum_{j=1}^{N} (\mathbf{Y}_j - \boldsymbol{\mu}_{\mathbf{Y}_j})' \Sigma_Y^{-1} (\mathbf{Y}_j - \boldsymbol{\mu}_{\mathbf{Y}_j}) \\ &= \text{Term}_1 + \sum_{j=1}^{N} (\mathbf{Y}_j - \boldsymbol{\mu}_{\mathbf{Y}_j})' B_1^{-1} \left( I_K \otimes \Sigma_E^{-1} - J_K \otimes W \right) B_1^{-1} (\mathbf{Y}_j - \boldsymbol{\mu}_{\mathbf{Y}_j}) \\ &= \text{Term}_1 + \sum_{j=1}^{N} \mathring{Y}_{\cdot j}' \left( I_K \otimes \Sigma_E^{-1} - J_K \otimes W \right) \mathring{Y}_{\cdot j} \\ &= \text{Term}_1 + \sum_{j=1}^{N} \mathring{Y}_{\cdot j}' \left( I_K \otimes \Sigma_E^{-1} \right) \mathring{Y}_{\cdot j} - \sum_{j=1}^{N} \mathring{Y}_{\cdot j}' \left( J_K \otimes W \right) \mathring{Y}_{\cdot j} \\ &= \text{Term}_1 + \text{Term}_2 - \text{Term}_3. \end{aligned} \tag{S1.25}$$

Here, $\mathring{Y}$ is the standardized data matrix defined in the following way

$$\mathring{Y} := B_1^{-1} \left( Y - \boldsymbol{\mu}_Y \right) = \underbrace{B_1^{-1} (Y - \mathbf{b}_0 \mathbf{1}_N')}_{Y^{(s)}} - \mathbf{1}_K \otimes (\beta X) \in \mathbb{R}^{mK \times N}. \tag{S1.26}$$

It is easy to see that $E\mathring{Y} = 0_{mK \times N}$, and

$$\text{cov}(\mathring{Y}_j) = B_1^{-1} \Sigma_Y B_1^{-1} = J_K \otimes (AA' + \Sigma_\Upsilon) + I_K \otimes \Sigma_E. \tag{S1.27}$$

For convenience, let us define sample-average of the original data as $\bar{Y} := \frac{1}{N} Y \cdot \mathbf{1}_N$; platform-average of the standardized data as $\bar{\mathring{Y}} := \frac{1}{K} \left( \mathbf{1}_K' \otimes I_m \right) \mathring{Y}$; and sample-average of the standardized data as

$$\breve{Y} := \frac{1}{N} \mathring{Y} \mathbf{1}_N = B_1^{-1} \left( \bar{Y} - \mathbf{b}_0 \right) - (\mathbf{1}_K \otimes \beta) \bar{X}. \tag{S1.28}$$

Apparently, $E\breve{Y} = \mathbf{0}_{mK}$, and $\text{cov}\left( \breve{Y} \right) = \frac{1}{N} \text{cov}(\mathring{Y}_j)$.

Using $\mathring{Y}$, the three terms in Equation (S1.25) can be expressed in

$$
\begin{aligned}
\mathrm{Term}_1 &:= mNK\log 2\pi + N\log\det(\Sigma_Y) \\
&= mNK\log(2\pi) + N\Big(\log C + 2\sum_{i,k}\log b_{1i,k} + (K-1)\sum_{i=1}^{m}\log\sigma_i^2 \\
&\quad + \sum_{i=1}^{m}\log(\sigma_i^2 + K\sigma_{\Upsilon,i}^2) + \sum_{l=1}^{L}\log(1+d_l^2)\Big). \\
\mathrm{Term}_2 &:= \sum_{j=1}^{N}(\mathbf{Y}_j - \boldsymbol{\mu}_{\mathbf{Y}_j})' B_1^{-1}\left(I_K\otimes\Sigma_E^{-1}\right)B_1^{-1}(\mathbf{Y}_j - \boldsymbol{\mu}_{\mathbf{Y}_j}) \\
&= \mathrm{tr}\left(\mathring{Y}'\left(I_K\otimes\Sigma_E^{-1}\right)\mathring{Y}\right) = \sum_{i,k}\sigma_i^{-2}\|\mathring{Y}_{i\cdot,k}\|^2. \\
\mathrm{Term}_3 &:= \sum_{j=1}^{N}(\mathbf{Y}_j - \boldsymbol{\mu}_{\mathbf{Y}_j})' B_1^{-1}\left(J_K\otimes W\right)B_1^{-1}(\mathbf{Y}_j - \boldsymbol{\mu}_{\mathbf{Y}_j}) \\
&= \mathrm{tr}\left(\mathring{Y}'\left(J_K\otimes W\right)\mathring{Y}\right) = K^2\cdot\mathrm{tr}\left(\bar{\mathring{Y}}'W\bar{\mathring{Y}}\right) \\
&= K^2\cdot\left[\sum_{i=1}^{m}\frac{\sigma_{\Upsilon,i}^2\|\bar{\mathring{Y}}_{i\cdot}\|^2}{\sigma_i^2(K\sigma_{\Upsilon,i}^2+\sigma_i^2)} + \mathrm{tr}\left(\bar{\mathring{Y}}'\tilde{U}M\tilde{U}'\bar{\mathring{Y}}\right)\right].
\end{aligned}
\tag{S1.29}
$$

Before we move on to the next section, we derive the total differentials of $\mathring{Y}$, $\bar{\mathring{Y}}$, and $\breve{Y}$, which will be useful in deriving gradient equations for conditional MLEs (maximum likelihood estimators).

$$
\begin{aligned}
\mathrm{d}\mathring{Y} &= \mathrm{d}(B_1^{-1})(Y - \mathbf{b}_0\mathbf{1}_N') - B_1^{-1}\mathrm{d}\mathbf{b}_0\mathbf{1}_N' - \mathbf{1}_K\otimes(\mathrm{d}\beta X). \\
\mathrm{d}\mathring{Y}_{i\cdot,k} &= \mathrm{d}(b_{1i,k}^{-1})(Y_{i\cdot,k} - b_{0i,k}\mathbf{1}_N') - b_{1i,k}^{-1}\cdot\mathrm{d}b_{0i,k}\cdot\mathbf{1}_N' - \mathrm{d}\beta_{i\cdot}X. \\
\mathrm{d}\bar{\mathring{Y}} &= \frac{1}{K}\left(\mathbf{1}_K'\otimes I_m\right)\left(\mathrm{d}(B_1^{-1})(Y - \mathbf{b}_0\mathbf{1}_N') - B_1^{-1}\mathrm{d}\mathbf{b}_0\mathbf{1}_N' - \mathbf{1}_K\otimes(\mathrm{d}\beta X)\right). \\
\left[\mathrm{d}\bar{\mathring{Y}}\right]_{ij} &= \frac{1}{K}\sum_{k=1}^{K}\left(\mathrm{d}(b_{1i,k}^{-1})(Y_{ij,k} - b_{0i,k}) - b_{1i,k}^{-1}\mathrm{d}b_{0i,k}\right) - [\mathrm{d}\beta X]_{ij}. \\
\mathrm{d}\breve{Y} &= \mathrm{d}(B_1^{-1})(\bar{Y} - \mathbf{b}_0) - B_1^{-1}\mathrm{d}\mathbf{b}_0 - \mathbf{1}_K\otimes(\mathrm{d}\beta\bar{X}).
\end{aligned}
\tag{S1.30}
$$

### S2 Parameter Estimation

#### S2.A Estimators of $\mathbf{b}_0$ and $\beta$

Notice that $\mathrm{Term}_1$ defined in Equation (S1.29) does not depend on $\mathbf{b}_0$ and $\beta$. If parameters other than $\mathbf{b}_0$ and $\beta$ (including $\mathbf{b}_1$, $A$, $\Sigma_E$, and $\Sigma_\Upsilon$) are all fixed, we can derive the total differential of the log-likelihood function as

$$
\begin{aligned}
\mathrm{d}\ell &= -\frac{1}{2}\mathrm{d}\mathrm{tr}\left(\mathring{Y}'\left(I_K \otimes \Sigma_E^{-1} - J_K \otimes W\right)\mathring{Y}\right) = -\mathrm{tr}\left(\mathring{Y}'\left(I_K \otimes \Sigma_E^{-1} - J_K \otimes W\right)\mathrm{d}\mathring{Y}\right) \\
&= \mathrm{tr}\left(\mathring{Y}'\left(I_K \otimes \Sigma_E^{-1} - J_K \otimes W\right)\left(B_1^{-1}\mathrm{d}\mathbf{b}_0\mathbf{1}'_N + \mathbf{1}_K \otimes (\mathrm{d}\beta X)\right)\right).
\end{aligned}
\tag{S2.31}
$$

In the above equation, the differential of the log-likelihood related to $\mathrm{d}\mathbf{b}_0$ is

$$
\mathrm{tr}\left(\mathring{Y}'\left(I_K \otimes \Sigma_E^{-1} - J_K \otimes W\right)B_1^{-1}\mathrm{d}\mathbf{b}_0\mathbf{1}'_N\right) = N\mathrm{tr}\left(\check{Y}'\left(I_K \otimes \Sigma_E^{-1} - J_K \otimes W\right)B_1^{-1}\mathrm{d}\mathbf{b}_0\right).
$$

Therefore

$$
\begin{aligned}
\nabla_{\mathbf{b}_0}\ell &= NB_1^{-1}\left(I_K \otimes \Sigma_E^{-1} - J_K \otimes W\right)\check{Y} \\
&= NB_1^{-1}\left(I_K \otimes \Sigma_E^{-1} - J_K \otimes W\right)\left[B_1^{-1}\left(\bar{Y} - \mathbf{b}_0\right) - (\mathbf{1}_K \otimes \beta)\bar{X}\right].
\end{aligned}
\tag{S2.32}
$$

It can be shown that the matrix $B_1^{-1}\left(I_K \otimes \Sigma_E^{-1} - J_K \otimes W\right)$ is always of full rank, so

$$
\nabla_{\mathbf{b}_0}\ell = 0 \iff B_1^{-1}\left(\bar{Y} - \mathbf{b}_0\right) - (\mathbf{1}_K \otimes \beta)\bar{X} = \mathbf{0} \Longrightarrow \hat{\mathbf{b}}_0 = \bar{Y} - B_1(\mathbf{1}_K \otimes \beta)\bar{X}. \tag{S2.33}
$$

Due to the identifiability constraint (S1.13b), we know that

$$
(\mathbf{1}_K \otimes \beta)\bar{X} = \mathrm{vec}\left(\beta\bar{X}\mathbf{1}_K^{\top}\right) = \mathbf{0}.
$$

Therefore the conditional Maximum Likelihood Estimate (MLE) of $\mathbf{b}_0$ is:

$$
\hat{\mathbf{b}}_0 = \bar{Y} = \frac{1}{N}Y\mathbf{1}_N. \tag{S2.34}
$$

On the other hand, in Equation (S2.31), terms related to $\mathrm{d}\beta$ is

$$
\begin{aligned}
&\mathrm{tr}\left(X\mathring{Y}'\left(I_K \otimes \Sigma_E^{-1} - J_K \otimes W\right)(\mathbf{1}_K \otimes \mathrm{d}\beta)\right) \\
&\quad = \mathrm{tr}\left(X\mathring{Y}'\left[\mathbf{1}_K \otimes \left((\Sigma_E^{-1} - KW)\mathrm{d}\beta\right)\right]\right) = K\mathrm{tr}\left(X\bar{\mathring{Y}}'(\Sigma_E^{-1} - KW)\mathrm{d}\beta\right).
\end{aligned}
\tag{S2.35}
$$

Therefore

$$
\nabla_{\beta}\ell = K(\Sigma_E^{-1} - KW)\bar{\mathring{Y}}X'. \tag{S2.36}
$$

It can be shown that $\Sigma_E^{-1} - KW$ is of full rank, therefore $\nabla_{\beta}\ell = \mathbf{0}$ iff $\bar{\mathring{Y}}X' = \mathbf{0}$. Define $\bar{\mathring{Y}}$ to be the sample mean of $\mathring{Y}$ taken over all technical platforms, i.e.

$$
\bar{\mathring{Y}} := \frac{1}{K}\left(\mathbf{1}'_K \otimes I_m\right)\mathring{Y}, \qquad \bar{\mathring{Y}}_{ij} = \frac{1}{K}\sum_{k=1}^{K}\mathring{Y}_{ij,k}. \tag{S2.37}
$$

Clearly, the covariance of $\bar{\mathring{Y}}_j$ is

$$
\mathrm{cov}\left(\bar{\mathring{Y}}_j\right) = AA' + \Sigma_{\Upsilon} + \frac{\Sigma_E}{K}. \tag{S2.38}
$$

Define

$$\tilde{Y} := \frac{1}{K}\left(\mathbf{1}'_K \otimes I_m\right) \underbrace{\left[B_1^{-1}\left(Y - \mathbf{b}_0 \mathbf{1}'_N\right)\right]}_{=Y^{(s)}}. \tag{S2.39}$$

By definition, $\tilde{Y}$ is the platform-mean of $Y^{(s)}$ defined in Equation (S1.26). Note that in order to compute $\tilde{Y}$, we need $\mathbf{b}_0$ and $\mathbf{b}_1$ but not $\beta$ and other parameters.

Because

$$\tilde{\mathring{Y}} = \frac{1}{K}\left(\mathbf{1}'_K \otimes I_m\right)\left[B_1^{-1}\left(Y - \mathbf{b}_0\mathbf{1}'_N\right) - (\mathbf{1}_K \otimes \beta)X\right] = \tilde{Y} - \beta X, \tag{S2.40}$$

we conclude that

$$\nabla_\beta \ell = \mathbf{0} \iff \tilde{Y}X' - \beta XX' = 0 \Longrightarrow \hat{\beta} := \tilde{Y}X'\left(XX'\right)^{-1}. \tag{S2.41}$$

As a remark, we note that $\hat{\mathbf{b}}_0$ defined in Equation (S2.34) and $\hat{\beta}$ defined in Equation (S2.41) are *conditional* MLE only because their *derivations* depend on fixing other parameters in the log-likelihood. That being said, these two estimators do not depend on covariance-related parameters: $A$, $\Sigma_E$, and $\Sigma_\Upsilon$.

#### S2.B Gradient of $\mathbf{b}_1$

Now let us focus on $\mathbf{b}_1$ and assume that all other parameters are fixed. We found it is easier to derive $\nabla_{\mathbf{b}_1}\ell$ from the total differential w.r.t. $\mathbf{b}_1^{-1}$ instead of $\mathbf{b}_1$, because it avoids awkward Jacobians.

Note that for an arbitrary differentiable function $\ell(\mathbf{b}_1)$, we have the following result that links the gradient of $\ell$ w.r.t $\mathbf{b}_1^{-1}$ and $\mathbf{b}_1^{-1}$

$$\nabla_{\mathbf{b}_1}\ell = \frac{D\mathbf{b}_1^{-1}}{D\mathbf{b}_1}\nabla_{\mathbf{b}_1^{-1}}\ell = \nabla_{\mathbf{b}_1^{-1}}\ell \circ \left(-\mathbf{b}_1^2\right). \tag{S2.42}$$

Using Equations (S1.25) and (S1.20), we obtain

$$\nabla_{\mathbf{b}_1^{-1}}\mathrm{Term}_1 = -2N\mathbf{b}_1, \qquad \nabla_{\mathbf{b}_1}\mathrm{Term}_1 = -2N\mathbf{b}_1 \circ \left(-\mathbf{b}_1^{-2}\right) = 2N\mathbf{b}_1^{-1}. \tag{S2.43}$$

The gradient of the rest terms in Equation (S1.25) w.r.t. $\mathbf{b}_1^{-1}$ is

$$\begin{aligned}
\mathrm{Rest} &:= \mathrm{Term}_2 - \mathrm{Term}_3 = \mathrm{tr}\left(\mathring{Y}'\left(I_K \otimes \Sigma_E^{-1} - J_K \otimes W\right)\mathring{Y}\right). \\
\mathrm{dRest} &= 2\mathrm{tr}\left(\mathring{Y}'\left(I_K \otimes \Sigma_E^{-1} - J_K \otimes W\right)\mathrm{d}(B_1^{-1})(Y - \mathbf{b}_0\mathbf{1}'_N)\right). \\
\nabla_{\mathbf{b}_1^{-1}}\mathrm{Rest} &= 2\mathrm{diag}\left((Y - \mathbf{b}_0\mathbf{1}'_N)\mathring{Y}'\left(I_K \otimes \Sigma_E^{-1} - J_K \otimes W\right)\right).
\end{aligned} \tag{S2.44}$$

Here $\mathrm{diag}(\cdot)$ extracts the diagonal of a square matrix as a vector.

While Equation (S2.44) provides a closed-form gradient of Rest w.r.t. $\mathbf{b}_1^{-1}$, it must be further simplified to be useful in practice because $(Y - \mathbf{b}_0\mathbf{1}'_N)\mathring{Y}'\left(I_K \otimes \Sigma_E^{-1} - J_K \otimes W\right)$ is a $mK \times mK$-dimensional matrix, which is prohibitively large to compute for real data.

We notice that $\mathring{Y}'\left(I_K \otimes \Sigma_E^{-1}\right)$ is just the transpose of per-row multiplication of $\mathring{Y}_j$ by $\sigma_i^{-2}$, which is computationally inexpensive.

It can be shown that $\mathring{Y}'\left(J_K \otimes W\right) = K(\mathbf{1}_K' \otimes F')$, where

$$F = W'\bar{\mathring{Y}} = \mathrm{Diag}\left(\frac{\sigma_{\Upsilon,i}^2}{\sigma_i^2(K\sigma_{\Upsilon,i}^2 + \sigma_i^2)}\right)\bar{\mathring{Y}} + (\tilde{U}M)(\tilde{U}'\bar{\mathring{Y}}). \tag{S2.45}$$

The first part of the above equation is a per-row multiplication applied to $\bar{\mathring{Y}} \in \mathbb{R}^{m\times N}$ therefore very efficient. The second part involves matrix multiplications of $\tilde{U} \in \mathbb{R}^{m\times L}$, $M \in \mathbb{R}^{L\times L}$, and $\bar{\mathring{Y}} \in \mathbb{R}^{m\times N}$, which are several magnitudes smaller than the naive computation defined in Equation (S2.44) for high-throughput data.

Using the above computational shortcut and notice that $\mathrm{diag}(\cdot)$ is the operator that extracts the diagonal elements of a square matrix, we can re-write $\nabla_{\mathbf{b}_1^{-1}}\mathrm{Rest}$ as

$$\nabla_{\mathbf{b}_1^{-1}}\mathrm{Rest} = 2\left[(Y - \mathbf{b}_0\mathbf{1}_N') \circ \mathrm{Second}\right]\mathbf{1}_N.$$
$$\mathrm{Second} := \left(I_K \otimes \Sigma_E^{-1} - J_K \otimes W\right)\mathring{Y} = (I_K \otimes \Sigma_E^{-1})\mathring{Y} - K(\mathbf{1}_K \otimes F).$$

Consequently

$$\begin{aligned}\nabla_{\mathbf{b}_1}\ell &= -\frac{1}{2}\left(\nabla_{\mathbf{b}_1}\mathrm{Term}_1 - \nabla_{\mathbf{b}_1^{-1}}\mathrm{Rest} \circ \mathbf{b}_1^{-2}\right) \\ &= -N\mathbf{b}_1^{-1} + \left(\left[(Y - \mathbf{b}_0\mathbf{1}_N') \circ \mathrm{Second}\right]\mathbf{1}_N\right) \circ \mathbf{b}_1^{-2}.\end{aligned} \tag{S2.46}$$

The above gradient leads to a highly nonlinear equation of $\mathbf{b}_1$ that does not have closed-form solutions. Instead, we will use a search algorithm (Algorithm 1) to find an approximate solution.

#### S2.C Gradient Descent for Estimating $\hat{\mathbf{b}}_1$

Below we describe a stable and scalable gradient descent procedure for estimating $\hat{\mathbf{b}}_1$. The steps are summarized in the algorithm below, followed by a derivation of the line search formula.

**Derivation of Line Search Step Size.** Let the univariate function of the step $s$

$$f^{(j)}(s) := \ell(\mathbf{b}_1^{(j)} + s\mathbf{v}^{(j)}). \tag{S2.47}$$

be the log-likelihood restricted in the line that passes through $\mathbf{b}_1^{(j)}$ and in the direction of $\nabla_{\mathbf{b}_1}^{(j)}$. Then,

$$\frac{\mathrm{d}f^{(j)}(0)}{\mathrm{d}s} = \|\nabla_{\mathbf{b}_1}^{(j)}\|, \qquad \frac{\mathrm{d}^2 f^{(j)}(0)}{\mathrm{d}s^2} = \left(\mathbf{v}^{(j)}\right)' H(\mathbf{b}_1^{(j)})\mathbf{v}^{(j)}, \tag{S2.48}$$

where $H(\mathbf{b}_1^{(j)}) \in \mathbb{R}^{(mK+1)\times(mK+1)}$ is the Hessian matrix of the log-likelihood function evaluated at $\mathbf{b}_1^{(j)}$. However, direct computation of $H(\mathbf{b}_1^{(j)})$ is infeasible for high-throughput data, so we propose to approximate it by a scalar matrix $H(\mathbf{b}_1^{(j)}) \approx h^{(j)} I_{mK+1}$, where

$$h^{(j)} := \frac{\langle \nabla_{\mathbf{b}_1}^{(j)} - \nabla_{\mathbf{b}_1}^{(j-1)},\ \mathbf{v}^{(j)}\rangle}{\langle s^{(j-1)}\mathbf{v}^{(j-1)},\ \mathbf{v}^{(j)}\rangle} = \frac{\|\nabla_{\mathbf{b}_1}^{(j)}\| - \|\nabla_{\mathbf{b}_1}^{(j-1)}\|\langle \mathbf{v}^{(j-1)},\ \mathbf{v}^{(j)}\rangle}{s^{(j-1)}\langle \mathbf{v}^{(j-1)},\ \mathbf{v}^{(j)}\rangle}. \tag{S2.49}$$

**Algorithm 1** Gradient Descent for $\mathbf{b}_1$

**Input:** Observed expression matrix $Y$, previously estimated $\mathbf{b}_0, \beta, \Sigma_E, \Sigma_\Upsilon$, initial $\mathbf{b}_1^{(0)}$, step size $s^{(0)} = 0.01$, tolerance `tol`, and maximum step size $s_{\max}$.

**Output:** Updated $\mathbf{b}_1$.

**Step 1: Initialization**

Compute $B_1^{(0)} = \text{Diag}(\mathbf{b}_1^{(0)}) \in \mathbb{R}^{mK \times mK}$, which is a block-diagonal matrix representing the platform-specific slopes.

To facilitate computation, standardize data across platforms, by removing platform-specific effects and scaling: $\mathring{Y} = (B_1^{(0)})^{-1}(Y - \mathbf{b}_0 \mathbf{1}_N') - \mathbf{1}_K \otimes (\beta X)$.

Compute the initial gradient $\nabla_{\mathbf{b}_1}^{(0)}$.

Compute the initial direction vector: $\mathbf{v}^{(0)} = \frac{\nabla_{\mathbf{b}_1}^{(0)}}{\|\nabla_{\mathbf{b}_1}^{(0)}\|}$.

Perform the first gradient descent step: $\mathbf{b}_1^{(1)} = \mathbf{b}_1^{(0)} + s^{(0)}\mathbf{v}^{(0)}$.

**Step 2: Iterative Updates**

**for** $j = 1, 2, \ldots$ **do** ▷ Until convergence criteria are met

Compute the updated gradient at the current step: $\nabla_{\mathbf{b}_1}^{(j)} =$ Gradient at $\mathbf{b}_1^{(j)}$.

Compute the new direction vector: $\mathbf{v}^{(j)} = \frac{\nabla_{\mathbf{b}_1}^{(j)}}{\|\nabla_{\mathbf{b}_1}^{(j)}\|}$.

Perform line search to adapt the step size $s^{(j)}$:

$$s^{(j)} = r^{(j)} s^{(j-1)}, \quad r^{(j)} = \frac{\|\nabla_{\mathbf{b}_1}^{(j)}\| \langle \mathbf{v}^{(j-1)}, \mathbf{v}^{(j)} \rangle}{\|\nabla_{\mathbf{b}_1}^{(j-1)}\| \langle \mathbf{v}^{(j-1)}, \mathbf{v}^{(j)} \rangle - \|\nabla_{\mathbf{b}_1}^{(j)}\|}.$$

If $s^{(j)} < 0$ or $s^{(j)} > s_{\max}$, set $s^{(j)} = s_{\max}$ to ensure numerical stability.

Update $\mathbf{b}_1$: $\mathbf{b}_1^{(j+1)} = \mathbf{b}_1^{(j)} + s^{(j)}\mathbf{v}^{(j)}$.

Check convergence:

**if** $|\ell(\mathbf{b}_1^{(j+1)}) - \ell(\mathbf{b}_1^{(j)})| < tol$ **then** ▷ Converged

**Break.**

**end if**

**end for**

**Return:** Final updated $\mathbf{b}_1$.

Using a second-order Taylor expansion,

$$f^{(j)}(s) \approx \ell(\mathbf{b}_1^{(j)}) + s\|\nabla_{\mathbf{b}_1}^{(j)}\| + \frac{s^2}{2} \cdot h^{(j)}, \qquad \frac{\mathrm{d}f^{(j)}(s)}{\mathrm{d}s} \approx \|\nabla_{\mathbf{b}_1}^{(j)}\| + h^{(j)}s. \tag{S2.50}$$

Therefore, the optimal step size is where the derivative $\frac{\mathrm{d}f^{(j)}(s)}{\mathrm{d}s}$ vanishes:

$$s^{(j)} := -\frac{\|\nabla_{\mathbf{b}_1}^{(j)}\|}{h^{(j)}} = r^{(j)}s^{(j-1)}, \qquad r^{(j)} := \frac{\|\nabla_{\mathbf{b}_1}^{(j)}\|\langle \mathbf{v}^{(j-1)},\ \mathbf{v}^{(j)}\rangle}{\|\nabla_{\mathbf{b}_1}^{(j-1)}\|\langle \mathbf{v}^{(j-1)},\ \mathbf{v}^{(j)}\rangle - \|\nabla_{\mathbf{b}_1}^{(j)}\|}.$$

This update ensures efficient progress while approximating second-order behavior without computing the full Hessian.

#### S2.D Conditional MLEs for Covariance-Related Parameters

In this section, we fix $\mathbf{b}_0$, $\mathbf{b}_1$, and $\beta$, and derive conditional MLEs for covariance-related parameters: $A$ ($U$ and $d_l^2$), $\Sigma_E$ ($\sigma_i^2$), and $\Sigma_\Upsilon$ ($\sigma_{\Upsilon,i}^2$).

To maximize the utility of the algorithm developed in this step, we first start with a latent factor model we call the Generalized Probabilistic Principal Component Analysis (GPPCA). Let $\mathbf{z} \sim N(0, I_L)$, $\mathbf{e} \sim N(\mathbf{0}_m, I_m)$, $\mathbf{u}, \boldsymbol{\tau}^2 \in \mathbb{R}^m$, $A \in \mathbb{R}^{m\times L}$, and

$$\mathring{\mathbf{y}} = A\mathbf{z} + \mathrm{Diag}(\boldsymbol{\tau})\mathbf{e}, \qquad \mathring{\mathbf{y}} \sim N\left(\mathbf{0}_m, \Sigma_{\mathring{\mathbf{y}}}\right), \quad \Sigma_{\mathring{\mathbf{y}}} = AA' + \mathrm{Diag}\left(\boldsymbol{\tau}^2\right). \tag{S2.51}$$

In the above GPPCA model, random vector $\mathring{\mathbf{y}} \in \mathbb{R}^m$ can be considered as $\mathring{Y}$ (defined in Equation (S1.26)) for one sample on one platform, with $\mathrm{Diag}\left(\boldsymbol{\tau}^2\right) = \Sigma_\Upsilon + \Sigma_E = \mathrm{Diag}(\sigma_{\Upsilon,i}^2 + \sigma_i^2)$.

##### S2.D.1 Estimate $AA'$

Let $W := \mathrm{Diag}(\boldsymbol{\tau}^{-1})$, it is easy to see that $W^2 = \mathrm{Diag}\left(\boldsymbol{\tau}^2\right)^{-1}$. Let $\tilde{A} := WA$, and $\tilde{A} = \tilde{U}\mathrm{Diag}\left(\tilde{d}_l\right)\tilde{V}'$ be its SVD. Using the Woodbury identity, we have

$$\begin{aligned}
\Sigma_{\mathring{\mathbf{y}}}^{-1} &= \mathrm{Diag}\left(\boldsymbol{\tau}^2\right)^{-1} - \mathrm{Diag}\left(\boldsymbol{\tau}^2\right)^{-1} A\left(I + A'\mathrm{Diag}\left(\boldsymbol{\tau}^2\right)^{-1} A\right)^{-1} A'\mathrm{Diag}\left(\boldsymbol{\tau}^2\right)^{-1} \\
&= W^2 - W^2A\left(I_L + A'W^2A\right)^{-1} A'W^2 = WPW. \\
P &:= I_m - \tilde{A}\left(I_L + \tilde{A}\tilde{A}'\right)^{-1}\tilde{A}' = I_m - \tilde{U}\mathrm{Diag}\left(\frac{\tilde{d}_l^2}{1+\tilde{d}_l^2}\right)\tilde{U}'.
\end{aligned} \tag{S2.52}$$

A useful fact is that

$$A'\Sigma_{\mathring{\mathbf{y}}}^{-1} = \tilde{A}'\left(I_m - \tilde{U}\mathrm{Diag}\left(\frac{\tilde{d}_l^2}{1+\tilde{d}_l^2}\right)\tilde{U}'\right)W = \tilde{V}\mathrm{Diag}\left(\frac{\tilde{d}_l}{1+\tilde{d}_l^2}\right)\tilde{U}'W.$$

Let $\mathring{Y}_k \in \mathbb{R}^{m\times N}$ be a sample of $N$ realizations of $\mathring{\mathbf{y}}$. It is easy to see that

$$-2\ell(A, \boldsymbol{\tau}^2|\mathring{Y}_k) = mN\log 2\pi + N\log|\Sigma_{\mathring{\mathbf{y}}}| + \mathrm{tr}\left(\mathring{Y}_k'\Sigma_{\mathring{\mathbf{y}}}^{-1}\mathring{Y}_k\right). \tag{S2.53}$$

Using matrix calculus, we know that

$$\mathrm{d}\Sigma_{\mathring{\mathbf{y}}} = \mathrm{d}AA' + A\mathrm{d}A' + \mathrm{Diag}\left(\mathrm{d}\boldsymbol{\tau}^2\right). \tag{S2.54}$$

$$\begin{aligned}\mathrm{d}\log|\Sigma_{\mathring{\mathbf{y}}}| &= \mathrm{tr}\left(\Sigma_{\mathring{\mathbf{y}}}^{-1}\mathrm{d}\Sigma_{\mathring{\mathbf{y}}}\right) = 2\mathrm{tr}\left(A'\Sigma_{\mathring{\mathbf{y}}}^{-1}\mathrm{d}A\right) + \mathrm{tr}\left(\Sigma_{\mathring{\mathbf{y}}}^{-1}\mathrm{Diag}\left(\mathrm{d}\boldsymbol{\tau}^2\right)\right) \\ &= 2\mathrm{tr}\left(\tilde{V}\mathrm{Diag}\left(\frac{\tilde{d}_l}{1+\tilde{d}_l^2}\right)\tilde{U}'W\mathrm{d}A\right) + \mathrm{tr}\left(WPW\mathrm{Diag}\left(\mathrm{d}\boldsymbol{\tau}^2\right)\right).\end{aligned} \tag{S2.55}$$

Now define $\tilde{Y}_k := W\mathring{Y}_k$, the scaled version of $\mathring{Y}_k$. Apparently

$$\Sigma_{\tilde{Y}_k} = WAA'W + I_m = \tilde{A}\tilde{A} + I_m. \tag{S2.56}$$

Let $H := \tilde{Y}_k\tilde{Y}_k'/N = W\mathring{Y}_k\mathring{Y}_k'W/N$, which is the MLE of $\Sigma_{\tilde{Y}_k}$. We have

$$\begin{aligned}\mathrm{dtr}\left(\mathring{Y}_k'\Sigma_{\mathring{\mathbf{y}}}^{-1}\mathring{Y}_k\right) &= -\mathrm{tr}\left(\Sigma_{\mathring{\mathbf{y}}}^{-1}\mathring{Y}_k\mathring{Y}_k'\Sigma_{\mathring{\mathbf{y}}}^{-1}\cdot\mathrm{d}\Sigma_{\mathring{\mathbf{y}}}\right) \\ &= -2\mathrm{tr}\left(A'\Sigma_{\mathring{\mathbf{y}}}^{-1}\mathring{Y}_k\mathring{Y}_k'\Sigma_{\mathring{\mathbf{y}}}^{-1}\mathrm{d}A\right) - \mathrm{tr}\left(\Sigma_{\mathring{\mathbf{y}}}^{-1}\mathring{Y}_k\mathring{Y}_k'\Sigma_{\mathring{\mathbf{y}}}^{-1}\cdot\mathrm{Diag}\left(\mathrm{d}\boldsymbol{\tau}^2\right)\right) \\ &= -2N\mathrm{tr}\left(\tilde{V}\mathrm{Diag}\left(\frac{\tilde{d}_l}{1+\tilde{d}_l^2}\right)\tilde{U}'HPW\mathrm{d}A\right) \\ &\quad - N\mathrm{tr}\left(WPHPW\mathrm{Diag}\left(\mathrm{d}\boldsymbol{\tau}^2\right)\right).\end{aligned} \tag{S2.57}$$

Therefore

$$\begin{aligned}-\frac{2}{N}\mathrm{d}\ell &= \mathrm{d}\log|\Sigma_{\mathring{\mathbf{y}}}| + \frac{1}{N}\mathrm{dtr}\left(\mathring{Y}_k'\Sigma_{\mathring{\mathbf{y}}}^{-1}\mathring{Y}_k\right) \\ &= 2\mathrm{tr}\left[\tilde{V}\mathrm{Diag}\left(\frac{\tilde{d}_l}{1+\tilde{d}_l^2}\right)\left(\tilde{U}' - \tilde{U}'HP\right)W\mathrm{d}A\right] \\ &\quad + \mathrm{tr}\left[W\left(P - PHP\right)W\mathrm{Diag}\left(\mathrm{d}\boldsymbol{\tau}^2\right)\right].\end{aligned} \tag{S2.58}$$

Because $\tilde{V}\mathrm{Diag}\left(\frac{\tilde{d}_l}{1+\tilde{d}_l^2}\right)$ and $W$ are invertible matrices, setting $\mathrm{d}\ell = 0$ for arbitrary $\mathrm{d}A$ leads to

$$\tilde{U}' - \tilde{U}'HP = \mathbf{0} \Longrightarrow PH\tilde{U} = \tilde{U}. \tag{S2.59}$$

Suppose $\tilde{U}$ is a set of eigenvector of $H$ such that $H\tilde{U} = \tilde{U}\Lambda$, where $\Lambda_{L\times L} = \mathrm{Diag}\left(\lambda_l\right)$. We have

$$\begin{aligned}PH\tilde{U} &= \left(I_m - \tilde{U}\mathrm{Diag}\left(\frac{\tilde{d}_l^2}{1+\tilde{d}_l^2}\right)\tilde{U}'\right)\tilde{U}\Lambda \\ &= \tilde{U}\left(\Lambda - \mathrm{Diag}\left(\frac{\tilde{d}_l^2}{1+\tilde{d}_l^2}\right)\Lambda\right) = \tilde{U}\mathrm{Diag}\left(\frac{\lambda_l}{1+\tilde{d}_l^2}\right).\end{aligned} \tag{S2.60}$$

Therefore, we only need to ensure that $1 + \tilde{d}_l^2 = \lambda_l$, or equivalently, $\tilde{d}_l^2 = \lambda_l - 1$. In summary, when $\boldsymbol{\tau}^2$ is given, we conclude that:

1. $\tilde{U}$ is the set of first $L$ eigenvectors of $H$, $\tilde{d}_l = \sqrt{\lambda_l - 1}$.

2. $\tilde{A}\tilde{A}' = \tilde{U}\text{Diag}\left(\lambda_l - 1\right)\tilde{U}'$, $AA' = U\text{Diag}\left(\lambda_l - 1\right)U'$, where $U := W^{-1}\tilde{U}$, $W^{-1} = \text{Diag}(\tau_i)$.

Note that $\tilde{V}$ is irrelevant because it is not part of the likelihood function (the likelihood function depends only on $AA'$).

Given the above results, we have

$$\tilde{A}\tilde{A}' = \tilde{U}\text{Diag}\left(\lambda_l - 1\right)\tilde{U}, \quad \frac{\tilde{d}_l^2}{1+\tilde{d}_l^2} = 1 - \frac{1}{\lambda_l}, \quad P = I_m - \tilde{U}\text{Diag}\left(1 - \frac{1}{\lambda_l}\right)\tilde{U}'.$$

$$\begin{aligned} PHP &= \left[I_m - \tilde{U}\text{Diag}\left(1 - \frac{1}{\lambda_l}\right)\tilde{U}'\right] H \left[I_m - \tilde{U}\text{Diag}\left(1 - \frac{1}{\lambda_l}\right)\tilde{U}'\right] \\ &= \left[H - \tilde{U}\text{Diag}(\lambda_l - 1)\tilde{U}'\right]\left[I_m - \tilde{U}\text{Diag}\left(1 - \frac{1}{\lambda_l}\right)\tilde{U}'\right] \\ &= H - \tilde{U}\text{Diag}\left(\lambda_l - 1\right)\tilde{U}' - \tilde{U}\text{Diag}(\lambda - 1)\tilde{U}' + \tilde{U}\text{Diag}\left(\lambda_l - 2 + \frac{1}{\lambda_l}\right)\tilde{U}' \\ &= H - \tilde{U}\text{Diag}\left(\frac{1-\lambda_l^2}{\lambda_l}\right)\tilde{U}'. \end{aligned}$$

$$P - PHP = I_m - H + \tilde{U}\text{Diag}\left(\lambda_l - 1\right)\tilde{U}' = I_m - H + \tilde{A}\tilde{A}'.$$

###### S2.D.2 Estimate $\tau_i^2$

Next, we derive an estimator of $\tau_i$ based on given $AA'$. By setting $\text{d}\ell = 0$ for arbitrary $\text{d}\boldsymbol{\tau}^2$ based on Equation (S2.58), we obtain $\text{diag}\left(P - PHP\right) = \mathbf{1}_m - \text{diag}(H) + \text{diag}(\tilde{A}\tilde{A}') = \mathbf{0}_m$.

By the definition of $H$, we know its diagonal elements are $\tau_i^{-2}\|\mathring{Y}_{k,i}\|^2/N$. Therefore $\boldsymbol{\tau}^2 \circ \text{diag}(H) = N^{-1}(\|\mathring{Y}_{k,1}\|^2, \ldots, \|\mathring{Y}_{k,m}\|^2)'$. Therefore

$$\boldsymbol{\tau}^2 \circ \left(\mathbf{1}_m - \text{diag}(H) + \text{diag}(\tilde{A}\tilde{A}')\right) = \boldsymbol{\tau}^2 - N^{-1}(\|\mathring{Y}_{k,1}\|^2, \ldots, \|\mathring{Y}_{k,m}\|^2)' + \text{diag}\left(AA'\right). \quad \text{(S2.61)}$$

Setting the above equation to zero, which is equivalent to setting the gradient of log-likelihood w.r.t. $\boldsymbol{\tau}^2$ to zero, we obtain the conditional MLE of $\tau_i$:

$$\hat{\tau}_i^2 = \frac{\|\mathring{Y}_{k,i}\|^2}{N} - \left(AA'\right)_{ii}. \quad \text{(S2.62)}$$

###### S2.D.3 Combined Parameter Estimation for GPPCA Model

Inspired by the above mathematical derivations, we propose the following algorithm to estimate $AA'$ and $\boldsymbol{\tau}^2$ from $\mathring{Y}_k$ for a GPPCA model. We assume that $L$, the number of singular values (PCs), is given.

1. **Data pre-processing**. For real data, we recommend: (a) center the data, and optionally, (b) standardize the data so that the sample STD of each $\mathring{\mathbf{y}}_i$ is constant one. Note that due to centering, we decide to replace $N$ by $N-1$ in some of the quantities used in theoretical derivations, e.g., we set $H := \tilde{Y}_k\tilde{Y}_k'/(N-1)$ instead of $\tilde{Y}_k\tilde{Y}_k'/N$.

2. **Initial estimation of $AA'$.** Conduct an initial PPCA on $\mathring{Y}_k$ (via SVD) to represent its covariance matrix as $\Sigma_{\mathring{\mathbf{y}}} = \widehat{AA'}^{(0)} + \tau^2 I_m$. Consider $\widehat{AA'}^{(0)}$ as the initial estimate of $AA'$. Specifically, let $\mathring{Y}_k = U_{\mathring{Y}_k} D_{\mathring{Y}_k} V'_{\mathring{Y}_k}$ be the SVD of $\mathring{Y}_k$. Let $U_{\mathring{Y}_k,L} \in \mathbb{R}^{m\times L}$ be the first $L$ columns (first $L$ left singular vectors) of $U_{\mathring{Y}_k}$, and $d_{\mathring{Y}_k,l}$ be the $l$th singular values in $D_{\mathring{Y}_k}$. It is easy to see that $U_{\mathring{Y}_k}$ is the eigenvector matrix of $\hat{\Sigma}_{\mathring{Y}_k} := \frac{\mathring{Y}_k \mathring{Y}'_k}{N-1}$ and the eigenvalues of $\hat{\Sigma}_{\mathring{Y}_k}$ (denoted by $\lambda_{\mathring{Y}_k,l}$) are related to $d_{\mathring{Y}_k,l}$ in this way: $\lambda_{\mathring{Y}_k,l} = \frac{d^2_{\mathring{Y}_k,l}}{N-1}$. We define the initial estimator of $AA'$ to be

$$\widehat{AA'}^{(0)} := U^{(0)}_{AA'} \mathrm{Diag}\left(d_l^{2,(0)}\right) U^{(0)'}_{AA'}. \qquad U^{(0)}_{AA'} := U_{\mathring{Y}_k,L}, \qquad d_l^{2,(0)} := \lambda_{\mathring{Y}_k,l} - \hat{\sigma}^{2,(0)}.$$
$$\hat{\tau}^{2,(0)} := \max\left(\frac{\sum_{l>L} \lambda_{\mathring{Y}_k,l}}{m-L}, \texttt{min.tau2}\right). \tag{S2.63}$$

Here `min.tau2` is a small preset positive number to prevent the under-estimation of the *i.i.d.* noise.

Note that $\widehat{AA'}^{(0)}$ defined in this way satisfies the following condition. Let $\hat{\Sigma}^{(0)}_{\mathring{Y}_k} := \widehat{AA'}^{(0)} + \hat{\tau}^{2,(0)} I_m$. We have

$$\mathrm{tr}\left(\hat{\Sigma}^{(0)}_{\mathring{Y}_k}\right) = \sum_{l=1}^{L} \lambda_{\mathring{Y}_k,l} - L\hat{\tau}^{2,(0)} + m\hat{\tau}^{2,(0)} = \sum_{l} \lambda_{\mathring{Y}_k,l} = \mathrm{tr}\left(\hat{\Sigma}_{\mathring{Y}_k}\right). \tag{S2.64}$$

For high-throughput data, $m$ can be much larger than $N$ and $L$. To save memory and computation time, we do not compute $\widehat{AA'}^{(0)}$ directly in this step. Instead, we save $U^{(0)}_{AA'}$ and $d_l^{2,(0)}$ for next steps. We also define $\hat{\boldsymbol{\tau}}^{2,(0)} := \hat{\tau}^{2,(0)} \mathbf{1}_m$, so that the quantity `err` can be computed in the first iteration (see below).

3. **Optional calculation related to standardization**. If the data are standardized, we know that $\Sigma_{\mathring{\mathbf{y}}}$ is also the *correlation* matrix of $\mathring{\mathbf{y}}$, meaning that $\mathrm{diag}(\Sigma_{\mathring{\mathbf{y}}}) = \mathbf{1}_m$. In this case, we standardize the estimates in this way

$$\begin{aligned}
\mathbf{s}^{(0)} &:= \mathrm{diag}\left(\widehat{AA'}^{(0)}\right) + \hat{\tau}^{2,(0)} \mathbf{1}_m = \left(U^{(0)}_{AA'} \circ U^{(0)}_{AA'}\right) \mathbf{d}^{2,(0)} + \hat{\tau}^{2,(0)} \mathbf{1}_m. \\
U^{(0s)}_{AA'} &:= \mathrm{Diag}\left(s_i^{(0)^{-1/2}}\right) U^{(0s)}_{AA'}, \qquad \hat{\boldsymbol{\tau}}^{2,(0s)} := \frac{\hat{\tau}^{2,(0)}}{\mathbf{s}^{(0)}}. \\
\widehat{AA'}^{(0s)} &:= \mathrm{Diag}\left(s_i^{(0)^{-1/2}}\right) \widehat{AA'}^{(0)} \mathrm{Diag}\left(s_i^{(0)^{-1/2}}\right) \\
&= U^{(0s)}_{AA'} \mathrm{Diag}\left(d_l^{2,(0)}\right) U^{(0s)'}_{AA'} + \mathrm{Diag}\left(\hat{\tau}_i^{2,(0s)}\right).
\end{aligned} \tag{S2.65}$$

It is easy to see that $\widehat{AA'}^{(0s)}$ thus computed is a correlation matrix. With a little abuse of the notation, we use simply denote the standardized estimates as $U^{(0)}_{AA'}$, $\hat{\boldsymbol{\tau}}^{2,(0)}$, and $\widehat{AA'}^{(0)}$, without the superscript "$s$". As an important remark, $U^{(0)}_{AA'}$ is no longer an *orthogonal matrix*. That being said, this will not hinder later computations.

When data are not standardized, we will need to compute the simple sample variance of $\mathring{Y}_k$, $\hat{\tau}^2_{\mathring{Y}_k,i} = \frac{\|\mathring{Y}_{k,i}\|^2}{N-1}$ (to be used in the iterations), before the iteration starts. If the data is standardized, we know that $\hat{\tau}^2_{\mathring{Y}_k,i} \equiv 1$ for all $i$.

4. **Update the estimate of $\boldsymbol{\tau}^2$**. This step is the start of an iteration. When $\widehat{AA'}^{(k-1)}$ is given in the form of $U^{(k-1)}_{AA'}$ and $\mathbf{d}^{2,(k-1)}$, we can compute the diagonal elements of $\widehat{AA'}^{(k-1)}$ efficiently in this way:

$$\text{diag}\left(\widehat{AA'}^{(k-1)}\right) = \left(U^{(k-1)}_{AA'} \circ U^{(k-1)}_{AA'}\right) \mathbf{d}^{2,(k-1)}. \tag{S2.66}$$

Note that this formula does not require $U^{(k-1)}_{AA'}$ to be orthogonal. We then use the following equation to estimate $\boldsymbol{\tau}^2$

$$\hat{\tau}_i^{2,(k)} := \max\left[\hat{\tau}^2_{\mathring{Y}_k,i} - \left(\widehat{AA'}^{(k-1)}\right)_{ii}, \texttt{min.tau2}\right]. \tag{S2.67}$$

Here `min.tau2` is a preset minimum value of $\hat{\tau}_i^{2,(k)}$. As a reminder, $\hat{\tau}^2_{\mathring{Y}_k,i} \equiv 1$ for the standardized data. Equation (S2.67) ensures that the resulting estimate of $\Sigma_{\mathring{\mathbf{y}}}$ is a correlation matrix.

5. **Update the estimate of $AA'$**. Given $\hat{\boldsymbol{\tau}}^{2,(k)}$, we define $\tilde{Y}^{(k)} := \text{Diag}(\hat{\boldsymbol{\tau}}^{-1,(k)})Y$ and conduct eigen-decomposition on $H^{(k)} := \frac{\tilde{Y}^{(k)}\tilde{Y}^{(k)\prime}}{N-1}$ to obtain the first $L$ eigenvectors (denoted by $U^{(k)}_{\tilde{Y},L}$) and eigenvalues (denoted by $\lambda^{(k)}_{\tilde{Y},l}$) via SVD on $\tilde{Y}^{(k)}$. We then compute the updated estimate of $AA'$, $\widehat{AA'}^{(k)} = U^{(k)}_{AA'}\text{Diag}\left(d_l^{2,(k)}\right) U^{(k)\prime}_{AA'}$, with the following formula

$$U^{(k)}_{AA'} := \text{Diag}(\hat{\boldsymbol{\tau}}^{1,(k)})U^{(k)}_{\tilde{Y},L}, \qquad d_l^{2,(k)} := \max\left(\lambda^{(k)}_{\tilde{Y},l} - 1,\, 0\right). \tag{S2.68}$$

6. **Stopping rule**. Repeat the iteration until the maximum number of iterations is reached or $\texttt{err}^{(k)} < \texttt{tol}$, where `tol` is a small preset threshold, and $\texttt{err}^{(k)}$ is a measure of the discrepancy between estimates computed in the $k$th and $(k-1)$th steps. Intuitively, we may want to define

$$\texttt{err}^{(k)} := \|\hat{\Sigma}^{(k)}_Y - \hat{\Sigma}^{(k-1)}_Y\|^2 = \|\widehat{AA'}^{(k)} - \widehat{AA'}^{(k-1)} + \text{Diag}(\hat{\boldsymbol{\tau}}^{2,(k)}) - \text{Diag}(\hat{\boldsymbol{\tau}}^{2,(k-1)})\|^2.$$

However, the above quantity requires the computation of very large matrices of dimension $m \times m$, which could take too much computational resources.

In practice, we define $\texttt{err}^{(k)}$ by the following equation, which can be computed efficiently

$$\begin{aligned}
\texttt{err}^{(k)} &:= \frac{1}{m^2}\left(\left\|\widehat{AA'}^{(k)} - \widehat{AA'}^{(k-1)}\right\|^2 + \left\|\hat{\boldsymbol{\tau}}^{2,(k+1)} - \hat{\boldsymbol{\tau}}^{2,(k)}\right\|^2\right) \\
m^2\texttt{err}^{(k)} &= \left\|\widehat{AA'}^{(k)}\right\|^2 + \left\|\widehat{AA'}^{(k-1)}\right\|^2 - 2\left\|\hat{A}^{(k)}\hat{A}^{(k-1)\prime}\right\|^2 + \left\|\hat{\boldsymbol{\tau}}^{2,(k+1)}\right\|^2 \\
&= \left\|B^{(k,k)}\right\|^2 + \left\|B^{(k-1,k-1)}\right\|^2 - 2\left\|B^{(k,k-1)}\right\|^2 \\
&\quad + \sum_{i=1}^{m}\left(\hat{\tau}_i^{2,(k+1)} - \hat{\tau}_i^{2,(k)}\right)^2.
\end{aligned} \tag{S2.69}$$

Here $B^{(k,k)}$, $B^{(k-1,k-1)}$, and $B^{(k,k-1)}$ are $(L\times L)$-dimensional matrices defined as follows

$$\begin{aligned}
B^{(k,k)} &:= \text{Diag}\left(\sqrt{d_l^{2,(k)}}\right) U_{AA'}^{(k)\prime} U_{AA'}^{(k)} \text{Diag}\left(\sqrt{d_l^{2,(k)}}\right), \\
B^{(k-1,k-1)} &:= \text{Diag}\left(\sqrt{d_l^{2,(k-1)}}\right) U_{AA'}^{(k-1)\prime} U_{AA'}^{(k-1)} \text{Diag}\left(\sqrt{d_l^{2,(k-1)}}\right), \\
B^{(k,k-1)} &:= \text{Diag}\left(\sqrt{d_l^{2,(k)}}\right) U_{AA'}^{(k)\prime} U_{AA'}^{(k-1)} \text{Diag}\left(\sqrt{d_l^{2,(k-1)}}\right).
\end{aligned} \tag{S2.70}$$

7. Denote the final iteration step by $k^*$. We consider $U_{AA'}^{(k^*)}$, $d_l^{2,(k^*)}$, and $\hat{\boldsymbol{\tau}}^{2,(k^*)}$ as the final results. Consequently, the final estimates of $AA'$ and $\Sigma_{\mathring{\mathbf{y}}}$ are defined as

$$\widehat{AA'}^{*} := U_{AA'}^{(k^*)} \text{Diag}\left(d_l^{2,(k^*)}\right) U_{AA'}^{(k^*)\prime}, \qquad \hat{\Sigma}_Y^* := \widehat{AA'}^{*} + \text{Diag}\left(\hat{\boldsymbol{\tau}}^{2,(k^*)}\right).$$

As a reminder, $U_{AA'}^{(k^*)}$ in general is not orthogonal due to variance re-scaling, but it does not affect the identifiability of $AA'$. However, this non-orthogonality may reduce the computational efficiency for downstream computations. Therefore, we define $\hat{A}^* := U_{AA'}^{(k^*)}\text{Diag}\left(\sqrt{d_l^{2,(k^*)}}\right)$, and apply SVD to $\hat{A}^*$ so that

$$\hat{A}^* = U^* D^* V^{*\prime}, \qquad \widehat{AA'}^{*} = \hat{A}^*\hat{A}^{*\prime} = U^* D^{2,*} U^{*\prime}. \tag{S2.71}$$

The final outputs of the algorithm are: $U^*$, $d_l^{2,*}$ (diagonal elements in $D^{2,*}$), and $\hat{\boldsymbol{\tau}}^{2,(k^*)}$. Due to numerical stability, we may need to run a last round of standardization of $\hat{A}^*$ and $\hat{\tau}_i^{2,(k^*)}$ to ensure that $\hat{\tau}_Y^* := \widehat{AA'}^{*} + \text{Diag}\left(\hat{\sigma}_i^{2,(k^*)}\right)$ is a correlation matrix.

#### S2.E Initial Estimation of Parameters

The performance of iterative numerical procedures, such as the gradient-descend method, always depends on a sensible initial parameter estimating method. We propose such a simple but effective parameter estimating procedure (Algorithm 2) to initialize the gradient descend method. This algorithm is implemented in function `InitEst()`.

**Algorithm 2** Initial Parameter Estimation for PXN

**Input:** Observed expression matrix $Y_k$ $(k = 1, \dots, K)$, covariate matrix $X$, and preset constants $w_i$, `min.sigma2`, `min.sigmaU2`.
**Output:** Initial estimates $\mathbf{b}_0^{(0)}, \mathbf{b}_1^{(0)}, \beta^{(0)}, \Sigma_E^{(0)}, \Sigma_\Upsilon^{(0)}$.

**Step 1: Compute Initial Intercept**
Center $X$, and compute the closed-form MLE for $\mathbf{b}_0^{(0)}$ as the sample mean of $Y$: $\mathbf{b}_0^{(0)}$ as the sample mean of $Y$: $\mathbf{b}_0^{(0)} = \frac{1}{N} Y \mathbf{1}_N$.

**Step 2: Compute Initial Slopes**
Perform an OLS regression of $Y$ on $X$ to remove deterministic terms and compute residuals: $R^{(0)} = Y - \beta^{(0)} X - \mathbf{b}_0^{(0)} \mathbf{1}'_N$.
Compute $\mathbf{b}_1^{(0)}$ as the empirical standard deviations of $R^{(0)}$.

**Step 3: Compute Initial Variance Estimates**
Standardize $R^{(0)}$ by removing $B_1^{(0)} = \mathrm{Diag}(\mathbf{b}_1^{(0)})$: $R^{(0,s)} = (B_1^{(0)})^{-1} R^{(0)}$.
Compute the average residual matrix $\bar{R}_j^{(0,s)}$ and estimate its covariance:

$$\bar{R}^{(0,s)} = \frac{1}{K}(1'_K \otimes I_m) R^{(0,s)}, \quad \mathrm{cov}(\bar{R}_j^{(0,s)}) \approx AA' + \Sigma_\Upsilon + \frac{1}{K}\Sigma_E.$$

**if** $K = 1$ **then** ▷ No pairing data
  **if** $L = 0$ **then** ▷ No latent factors
    The covariance simplifies to: $\mathrm{cov}(R_j^{(0,s)}) \approx \Sigma_\Upsilon + \Sigma_E = I_m$.
    Estimate: $\hat{\sigma}^2_{\Upsilon,i} = w_i$ and $\hat{\sigma}_i^2 = 1 - w_i$.
  **else** ▷ $L > 0$: Latent factors exist
    Use GPPCA (Section S2.D) to estimate $AA'$ (in terms of $\hat{U}_{AA'}, \hat{d}_l^2$) and $\hat{\tau}_i^2 \approx \sigma^2_{\Upsilon,i} + \sigma_i^2$.
    Estimate: $\hat{\sigma}^2_{\Upsilon,i} = w_i \cdot \hat{\tau}_i^2$ and $\hat{\sigma}_i^2 = (1 - w_i) \cdot \hat{\tau}_i^2$.
  **end if**
**else** ▷ $K > 1$: Pairing data exists
  Use GPPCA (Section S2.D) to estimate: $\hat{U}_{AA'}, \hat{d}_l^2$, and $\hat{\tau}_i^2 \approx \sigma^2_{\Upsilon,i} + \sigma_i^2/K$.
  Compute $\hat{\sigma}_i^2$ and $\hat{\sigma}^2_{\Upsilon,i}$:

$$\hat{\sigma}_i^2 = \max\left[\frac{K}{K-1}\left(1 - \mathrm{var}(\bar{R}_j^{(0,s)})\right),\ \texttt{min.sigma2}\right],$$

$$\hat{\sigma}^2_{\Upsilon,i} = \max\left[\hat{\tau}_i^2 - \frac{\hat{\sigma}_i^2}{K},\ \texttt{min.sigmaU2}\right].$$

**end if**

**Step 4: Compute Initial Regression Coefficients**
Compute $\beta^{(0)}$ using:

$$\beta^{(0)} = \tilde{Y} X'(XX')^{-1}, \quad \tilde{Y} = \frac{1}{K}(1'_K \otimes I_m)\left[(B_1^{(0)})^{-1}(Y - \mathbf{b}_0^{(0)} \mathbf{1}'_N)\right].$$



**Return:** $\mathbf{b}_0^{(0)}, \mathbf{b}_1^{(0)}, \beta^{(0)}, \Sigma_E^{(0)}, \Sigma_\Upsilon^{(0)}$.

We deliberately implement a special case with $K = 1$ (a single technical platform, i.e., no pairing data) in the function `InitEst()`. Although this setting is not relevant for cross-platform normalization, its inclusion allows users to fit a one-platform PXN model, which can be viewed as a computationally efficient probabilistic PCA (PPCA) model with gene-specific *i.i.d.* noise variances. We think this special case is still useful, because it enables a range of downstream analyses beyond normalization. For example, the fitted PXN model permits decomposition of gene expression into a latent factor component and a residual component. Because the model provides an estimate of the total *i.i.d.* variance ($\tau_i^2$) for each gene, it can be used for model-based outlier detection. In principle, this approach should be more accurate than methods based solely on marginal sample variances, as our approach accounts for inter-gene correlation through latent factors and incorporates covariate effects via the $\beta X$ component in the PXN model. Similarly, missing-value imputation based on the PXN model is expected to outperform marginal mean or median imputation by borrowing information efficiently across genes.

However, when only a single platform is available, the intrinsic biological variation $\sigma^2_{\Upsilon,i}$ cannot be separately identified from the measurement error variance $\sigma_i^2$; only their sum $\tau_i^2 = \sigma^2_{\Upsilon,i} + \sigma_i^2$ is identifiable. For several applications, including PXN-based outlier detection and missing-value imputation aforementioned, this decomposition is not strictly required. Nevertheless, to ensure consistent output of the function `InitEst()` across different choices of $K$, we introduce a set of user-specifiable weights $w_i \in (0, 1)$, $i = 1, \ldots, m$, which control the proportion of $\tau_i^2$ attributed to $\sigma^2_{\Upsilon,i}$ in the $K = 1$ case (see the "If $K = 1$" branch in Algorithm 2). By default, we set $w_i \equiv 0.5$, corresponding to an equal split between intrinsic variation and measurement error, while allowing experienced users to specify alternative values. We emphasize that in the typical setting of PXN with paired data ($K \geq 2$), $\sigma^2_{\Upsilon,i}$ and $\sigma_i^2$ are **identifiable** and are estimated directly without requiring user-specified weights $w_i$.

### S3 Model-based Cross-platform Normalization Procedures

This section provides technical details for the PXN procedure introduced in *Model-Based Cross-Platform Normalization* section of the main text.

Let $\mathbf{y}_k$ and $\mathbf{y}_{k'}$ be random vectors that represent gene expression profiles of a single biological sample quantified on the source platform $k$ and target platform $k'$, respectively. PXN uses the conditional expectation

$$\hat{\mathbf{y}}_{k' \leftarrow k} := E\big(\mathbf{y}_{k'} \mid \mathbf{y}_k, X, \hat{\theta}\big) = \boldsymbol{\mu}_{k'} + \Sigma_{k',k}\Sigma_k^{-1}\left(\mathbf{y}_k - \boldsymbol{\mu}_k\right). \tag{S3.72}$$

to predict $\mathbf{y}_{k'}$ (assuming unobserved) from the observed values $\mathbf{y}_k$. This predictor leverages clinical covariates $X$ and model parameters $\hat{\theta}$ estimated from the training data, and is the empirical best linear unbiased predictor (EBLUP) that enjoys many theoretical optimal properties based on our modeling assumptions.

For each platform $k = 1, \dots, K$, define $B_{1,k} := \text{Diag}(\mathbf{b}_{1,k})$. The model specifies:

$$\begin{aligned}
\boldsymbol{\mu}_k &= E(\mathbf{y}_k | X, \hat{\theta}) = B_{1,k}\beta X + \mathbf{b}_{0,k}1'_N, \\
\Sigma_k &= \text{Cov}(\mathbf{y}_k) = B_{1,k}(AA' + \Sigma_\Upsilon + \Sigma_E)B_{1,k}, \\
\Sigma_{k',k} &= \text{Cov}(\mathbf{y}_{k'}, \mathbf{y}_k) = B_{1,k'}(AA' + \Sigma_\Upsilon)B_{1,k}.
\end{aligned} \tag{S3.73}$$

Based on the Woodbury identity and the fact that $\Sigma_\Upsilon^{-1/2}AA'\Sigma_\Upsilon^{-1/2} = \tilde{A}\tilde{A}' = UD^2U$:

$$\begin{aligned}
(AA' + \Sigma_\Upsilon + \Sigma_E)^{-1} &= \Sigma_\Upsilon^{-1/2}\left(UD^2U' + I_m + \Sigma_\Upsilon^{-1}\Sigma_E\right)^{-1}\Sigma_\Upsilon^{-1/2} \\
&= \Sigma_\Upsilon^{-1/2}W_\Upsilon\left(I_m + \Sigma_\Upsilon^{-1}\Sigma_E - U\left(D^{-2} + U'W_\Upsilon U\right)^{-1}U'\right)W_\Upsilon\Sigma_\Upsilon^{-1/2}. \\
W_\Upsilon &:= \text{Diag}(w_{\Upsilon,i}) = \text{Diag}\left(\frac{\sigma^2_{\Upsilon,i}}{\sigma^2_{\Upsilon,i} + \sigma^2_i}\right).
\end{aligned}$$

Therefore

$$\begin{aligned}
\Sigma_k^{-1} &= B_{1,k}^{-1}\Sigma_\Upsilon^{-1/2}W_\Upsilon\left(I_m + \Sigma_\Upsilon^{-1}\Sigma_E - U\left(D^{-2} + U'W_\Upsilon U\right)^{-1}U'\right)W_\Upsilon\Sigma_\Upsilon^{-1/2}B_{1,k}^{-1} \\
&= B_{1,k}^{-1}\Sigma_\Upsilon^{-1/2}\left[W_\Upsilon - W_\Upsilon U\left(D^{-2} + U'W_\Upsilon U\right)^{-1}U'W_\Upsilon\right]\Sigma_\Upsilon^{-1/2}B_{1,k}^{-1}.
\end{aligned} \tag{S3.74}$$

And

$$\begin{aligned}
\Sigma_{k',k}\Sigma_k^{-1} &= B_{1,k'}\Sigma_\Upsilon^{1/2}\left(UD^2U' + I_m\right)\left[W_\Upsilon - W_\Upsilon U\left(D^{-2} + U'W_\Upsilon U\right)^{-1}U'W_\Upsilon\right]\Sigma_\Upsilon^{-1/2}B_{1,k}^{-1} \\
&= W_\Upsilon B_{1,k'}B_{1,k}^{-1} + B_{1,k'}M^{(2)}B_{1,k}^{-1}. \\
M^{(2)} &:= \Sigma_\Upsilon^{1/2}\left[UD^2 - (UD^2U' + I_m)W_\Upsilon U\left(D^{-2} + U'W_\Upsilon U\right)^{-1}\right]U'W_\Upsilon\Sigma_\Upsilon^{-1/2}.
\end{aligned} \tag{S3.75}$$

By combining Equations (S3.72) and (S3.75), we have the formula for cross-platform normalization (using $\mathbf{y}_k$ from platform $k$ to predict their equivalent values on platform $k'$):

$$\begin{aligned}
\hat{\mathbf{y}}_{k'\leftarrow k} &:= E\left(\mathbf{y}_{k'}|\mathbf{y}_k, X, \hat{\theta}\right) = \boldsymbol{\mu}_{k'} + \left[W_\Upsilon B_{1,k'}B_{1,k}^{-1} + B_{1,k'}M^{(2)}B_{1,k}^{-1}\right](\mathbf{y}_k - \boldsymbol{\mu}_k) \\
&= \boldsymbol{\mu}_{k'} + \text{GSI} + \text{SI}. \\
\text{GSI} &:= W_\Upsilon B_{1,k'}B_{1,k}^{-1}(\mathbf{y}_k - \boldsymbol{\mu}_k) = \text{Diag}\left(\frac{\sigma^2_{\Upsilon,i}}{\sigma^2_{\Upsilon,i} + \sigma^2_i}\right)B_{1,k'}\mathring{\mathbf{y}}_k. \\
\text{SI} &:= B_{1,k'}M^{(2)}B_{1,k}^{-1}(\mathbf{y}_k - \boldsymbol{\mu}_k) = B_{1,k'}M^{(2)}\mathring{\mathbf{y}}_k.
\end{aligned} \tag{S3.76}$$

In the above equation, GSI stands for the "Gene-Specific Information" which adjusts for platform-specific translation, scaling, and measurement noise:

$$\text{GSI}_i = \frac{\sigma^2_{\Upsilon,i}}{\sigma^2_{\Upsilon,i} + \sigma^2_i} \cdot \frac{b_{1i,k'}}{b_{1i,k}} \cdot (\mathbf{y}_{i,k} - \mu_{i,k}).$$

SI stands for the "Shared Information", which is a low-rank multivariate correction based on latent structure learned via PPCA. It can be understood heuristically as follows

$$\underbrace{B_{1,k'}\Sigma_\Upsilon^{1/2}\left[UD^2 - (UD^2U' + I_m)W_\Upsilon U\left(D^{-2} + U'W_\Upsilon U\right)^{-1}\right]}_{\text{turns PCs to enhanced prediction for } \mathbf{y}_{k'}} \times \underbrace{U'W_\Upsilon\Sigma_\Upsilon^{-1/2}\mathring{\mathbf{y}}_k}_{\text{top PCs of scaled } \mathring{\mathbf{y}}_k} \tag{S3.77}$$

To improve computational efficiency and modularity in the R implementation, we separate the computation of SI into the following steps.

1. Define `Term1` $:= \mathrm{diag}(B_{1,k'}\Sigma_\Upsilon^{1/2})$ and `PCs` $:= U'W_\Upsilon\Sigma_\Upsilon^{-1/2}\mathring{\mathbf{y}}_k$.

2. Define `UD2` $:= UD^2$, and `Term2` $:= W_\Upsilon U\left(D^{-2}+U'W_\Upsilon U\right)^{-1}$.

3. The final answer is

$$\mathrm{SI} = \texttt{Term1}(\texttt{UD2}\% * \%((I_L - U'\% * \%\texttt{Term2})\% * \%\texttt{PCs}) - \texttt{Term2}\% * \%\texttt{PCs}).$$

### S4 Model Bridging

#### S4.A Motivation

In real-world applications, the target platform is not always directly linked to all other platforms. Figure 1 in the main manuscript illustrates an example in which platform groups $\{A, B\}$ ($n_1 = 60$ samples) and $\{B, C, D\}$ ($n_2 = 80$ samples) are partially paired through a common platform $B$. Traditional pairwise normalization methods would have to rely on a sequence of "jumps" to perform indirect normalization. For example, normalizing data from platform $A$ to $B$, and then from $B$ to $C$. Such procedures are awkward, order-dependent, and prone to error accumulation. Moreover, another platform group $\{E, F\}$ ($n_3 = 100$ samples) is completely disconnected from the first four platforms, making indirect normalization impossible under pairwise methods.

From a theoretical perspective, even when samples are not paired across platform groups, the underlying **inter-gene correlation structure** should still be shared across all platforms. In other words, estimation of latent biological structure (via $AA'$) and intrinsic biological variation (via $\Sigma_\Upsilon$) should benefit from integrating data across disconnected platform groups.

The PXN framework eliminates the need for sequential jumps by enabling model-level bridging. Specifically, PXN assumes that all observed expression matrices $\{Y_k\}$ are platform-specific transformations of a common latent standardized expression $Y^{(0)}$, which captures covariate effects ($\beta X$), low-rank latent factors ($A\mathbf{Z}$), and intrinsic biological variation ($\Upsilon$). As a result, $Y^{(0)}$ provides a natural bridge linking all technical platforms, even in the absence of direct or indirect pairings.

#### S4.B Integrated PXN Estimation

Suppose the full set of platforms is partitioned into $g = 1, \ldots, G$ disjoint pairing groups. We first estimate PXN model parameters separately within each fully connected pairing group to obtain estimates $\hat{\theta}^{(g)}$ for $g = 1, \ldots, G$. We then combine these estimates to form a bridging PXN model, denoted by $\hat{\theta}^I$, which leverages all available training data and enables direct harmonization between any pair of platforms without jumping.

Due to technical reasons, different model parameters must be bridged using strategies tailored to their statistical properties, described as follows.

**Shared parameters $\beta$, $\Sigma_E$, and $\Sigma_\Upsilon$.** For these parameters shared across all platforms, we compute sample-size–weighted averages as the corresponding parameters for the bridged model:

$$\hat{\sigma}_i^{I,2} := \frac{1}{N}\sum_{g=1}^{G} n_g \hat{\sigma}_i^{2,(g)}, \quad \hat{\sigma}_{\Upsilon,i}^{I,2} := \frac{1}{N}\sum_{g=1}^{G} n_g \hat{\sigma}_{\Upsilon,i}^{2,(g)}, \quad \hat{\beta}^I := \frac{1}{N}\sum_{g=1}^{G} n_g \hat{\beta}^{(g)}, \tag{S4.78}$$

where $n_g$ is the sample size of the $g$th pairing group and $N = \sum_{g=1}^{G} n_g$.

**Latent factor covariance $AA'$.** Direct averaging of $\widehat{AA'}^{(g)}$ is computationally infeasible and numerically unstable. Instead, we estimate the integrated latent structure using pooled standardized residuals:

1. For each group $g$, compute standardized residuals $\mathring{Y}^{(g)}$ (see Equation (S1.26)) and their within-group platform means

$$\bar{\mathring{Y}}^{(g)} := \frac{1}{K_g}(\mathbf{1}'_{K_g} \otimes I_m)\mathring{Y}^{(g)} \in \mathbb{R}^{m \times n_g}.$$

2. Concatenate $\bar{\mathring{Y}}^{(g)}$ column-wise across all groups to form $\bar{\mathring{\mathbf{Y}}} \in \mathbb{R}^{m \times N}$. Perform SVD

$$\bar{\mathring{\mathbf{Y}}} = U_{\bar{\mathring{\mathbf{Y}}}} D_{\bar{\mathring{\mathbf{Y}}}} V'_{\bar{\mathring{\mathbf{Y}}}}.$$

   Let $U_L^I$ be the first $L$ columns of $U_{\bar{\mathring{\mathbf{Y}}}}$, and let $d_l^I$ be the corresponding singular values. The integrated estimate of $AA'$ is

$$\widehat{AA'}^I = U_L^I \mathrm{Diag}\big((d_l^I)^2\big)\, U_L^{I\prime}.$$

**Platform-specific parameters $\mathbf{b}_{0,k}$ and $\mathbf{b}_{1,k}$.** For platform-specific parameters $\mathbf{b}_{0,k}$ and $\mathbf{b}_{1,k}$, we average estimates across only those pairing groups in which platform $k$ appears:

$$\hat{\mathbf{b}}_{0,k}^I := \frac{\sum_{g\in\mathcal{S}_k} n_g \hat{\mathbf{b}}_{0,k}^{(g)}}{\sum_{g\in\mathcal{S}_k} n_g}, \qquad \hat{\mathbf{b}}_{1,k}^I := \frac{\sum_{g\in\mathcal{S}_k} n_g \hat{\mathbf{b}}_{1,k}^{(g)}}{\sum_{g\in\mathcal{S}_k} n_g}, \tag{S4.79}$$

where $\mathcal{S}_k$ denotes the set of pairing groups containing platform $k$.

Finally, we want to point out that we explored several alternative strategies for integrating latent structure estimates, but found them unsuitable for high-throughput data. Directly averaging $\widehat{AA'}^{(g)}$ across groups is computationally prohibitive due to the $m \times m$ matrix size and does not guarantee positive semi-definiteness, especially when the sample size is limited such that the uncertainly of $\widehat{AA'}^{(g)}$ is high. We also tried geometry-aware approaches, such as the generalized Procrustes analyses implemented in R package `shape`. While these procedures correctly preserve positive semi-definiteness, they are one to two orders of magnitude slower than the (already inefficient) direct averages, and require excessive memory, rendering them impractical for large-scale genomic applications.

### S5 Technical Details of Data Simulation

To evaluate the performance of PXN and compare it with existing normalization methods, we simulated a dataset using the generative model described in Equation (S1.11). This simulated data consisted of 1,000 gene expression profiles across $N = 240$ subjects sequenced on six technical platforms, resulting in a $6,000 \times 240$ matrix. From this full dataset, we extracted three subsets for subsequent analyses. Subset AB was a $2,000 \times 60$ matrix corresponding to cells A1 and B1, representing gene expressions from $n_1 = 60$ subjects measured on platforms A and B. Similarly, subset BCD was a $3,000 \times 80$ matrix consisted of cells B2, C2, and D2; and the subset EF was a $2,000 \times 100$ matrix composed of cells E3 and F3. Note that the white cells (e.g., A2, B3) corresponded to unobserved platforms thus were not included in the above three subsets. They were nonetheless generated in the overall simulation data and later used for evaluating the accuracy of cross-platform bridging procedures.

We constructed the design matrix $X$ with $p = 3$ covariates. The first covariate, $x_1$, was a binary variable representing the treatment assignment. The control group included the first 20, 50, and 30 subjects in subsets AB, BCD, and EF, respectively. Consequently, the treatment group consisted of the remaining 40, 30, and 70 subjects in the corresponding subsets. The second covariate, $x_2$, was an integer-valued random variable representing age, sampled uniformly from 18 to 80. The third covariate, $x_3$, was sampled from a standard normal distribution $N(0, 1)$, representing a confounding covariate that was not of primary interest in the differential expression (DE) analysis but needed to be considered for its potential confounding effects.

We then generated the corresponding gene-specific linear coefficients for gene $i$, denoted by $\beta_{i,\text{Treatment}}$, $\beta_{i,\text{Age}}$, and $\beta_{i,x_3}$. The first 100 genes were designated as treatment-associated DEGs, with $\beta_{i,\text{Treatment}} \sim U(0.5, 1.5)$ and $\beta_{i,\text{Treatment}} = 0$ for all other genes. The next 100 genes were DEGs associated with age, with $\beta_{i,\text{Age}} \sim U(-0.1, 0.1)$, and $\beta_{i,\text{Age}} = 0$ otherwise. For all genes, the confounder coefficient $\beta_{i,x_3}$ was sampled from $N(0, 1)$.

We generated platform- and gene-specific intercepts and slopes for $K = 6$ platforms, sampling each from *i.i.d.* normal distributions: $b_{0,i,k} \sim N(\mu_{b_0,k}, 0.5^2)$ and $b_{1,i,k} \sim N(\mu_{b_1,k}, 0.05^2)$. Platform-specific means were set as follows: $\mu_{b_0,A} = 8$, $\mu_{b_0,B} = 7.5$, $\mu_{b_0,C} = 7$, $\mu_{b_0,D} = 6.5$, $\mu_{b_0,E} = 6$, and $\mu_{b_0,F} = 5$. Gene gene-specific means were set as follows: $\mu_{b_1,A} = 0.5$, $\mu_{b_1,B} = 0.6$, $\mu_{b_1,C} = 0.55$, $\mu_{b_1,D} = 0.65$, $\mu_{b_1,E} = 0.45$, and $\mu_{b_1,F} = 0.7$. To ensure positive correlation across platforms, any $b_{1,i,k} < 0.1$ was replaced by 0.1.

The between-gene correlation structure used in the simulation studies was estimated from the real data used by Zhang et al. [2020]. Specifically, we took 1,000 randomly selected genes and estimated the correlation structure based on the PPCA model. The estimated number of PCs was $L = 8$, mean $\sigma_i^2 = 0.270$, and mean $\sigma_{\Upsilon,i}^2 = 0.424$.

### S6 Technical Details of Real Data Demonstration

#### S6.A Data Preparation of CellMiner Data

We evaluated PXN on the CellMiner molecular profiling resource from the U.S. National Cancer Institute, which provides multi-platform transcriptomic measurements for the NCI-

60 panel of human cancer cell lines. This dataset consists of data generated on six technical platforms. Specifically, it includes four generations of Affymetrix GeneChip platforms: Human Genome U95 (HG-U95 A–E; five-chip set; 65,000 probe sets), Human Genome U133 (HG-U133 A–B; two-chip set; 44,000 probe sets), Human Genome U133 Plus 2.0 ( 47,000 transcripts), and the exon-level Human Exon 1.0 ST array. In addition, CellMiner provides an Agilent Whole Human Genome Microarray (4×44K) and an RNA-seq dataset generated on Illumina HiSeq 2000 platform. All data were processed using standard summarization pipelines: the Affymetrix arrays were summarized using GCRMA, RMA, or MAS5; the Agilent data were processed via GeneSpring; and the RNA-seq reads were aligned to the hg19 reference genome using STAR with accompanying quality-control summaries. For readability, original CellMiner platform names were mapped to short labels (A = HG-U133, B = HG-U133 Plus 2.0, C = HG-U95, D = HuEx-1.0-ST, E = Agilent, F = RNA-seq).

Processed expression tables for each CellMiner platform were downloaded from the official CellMiner website. We restricted analyses to the intersection of samples observed across the six platforms. Concretely, we extracted the platform-specific sample columns, computed the set intersection, and reordered each platform's expression matrix to the same sample order. This ensured that (a) the same fold assignments could be reused across all platforms and methods and (b) comparisons across models were made on exactly the same held-out samples within each cross-validation fold.

Microarray platforms contain multiple probes or probesets mapping to the same gene, which induces duplicated gene identifiers and complicates cross-platform comparisons. Within each platform, we converted probe/probeset-level measurements to a gene-level matrix indexed by Entrez Gene ID. When multiple probes mapped to the same Entrez ID, we collapsed them by taking the sample-wise median across probes. For Affymetrix platforms, we additionally excluded `_x_at` probesets prior to collapsing because they are more prone to cross-hybridization. After collapsing, each platform contained at most one row per Entrez ID and no missing expression values.

After probe collapsing and sample alignment, each platform was represented as a numeric matrix with genes (Entrez IDs) as rows and shared samples as columns. For readability in the manuscript, original CellMiner platform names were mapped to short labels $\{A, B, C, D, E, F\}$.

**Cross-platform comparability** PXN and comparator methods were evaluated on feature spaces that match the genes used to fit each model. To support this, we precomputed intersections of Entrez IDs across platform subsets (Table S1). These intersections were then used to (i) define the training feature space for each model and (ii) ensure that MSPE was computed on the same genes used for that model. There is significant variation in the number of genes measured by each platform, ranging from 8,590 (HG-U95) to 28,287 (HuEx-1.0-ST). The Jaccard index was calculated to measure the similarity between any two gene sets by dividing the intersection of the sets by their union. Higher Jaccard means more similar gene coverage (e.g., HG-U133 Plus 2.0 vs RNA-seq = 0.780; HuEx-1.0-st vs HG-U95 = 0.302). Thus, though some are all Affymetrix platforms, they are not interchangeable in gene coverage. A subset of 7,441 genes was identified as being measured across all six platforms. This common gene set serves as the primary basis for direct gene-level cross-platform comparisons.

| Platform 1 | Platform 2 | $n_1$ | $n_2$ | $n_{\text{intersect}}$ | $n_{\text{union}}$ | Jaccard |
|---|---|---|---|---|---|---|
| HG-U133 (A) | HG-U133 Plus 2.0 (B) | 12514 | 20202 | 12513 | 20203 | 0.619 |
| HG-U133 (A) | HG-U95 (C) | 12514 | 8590 | 8476 | 12628 | 0.671 |
| HG-U133 (A) | HuEx-1.0-ST (D) | 12514 | 28287 | 12377 | 28424 | 0.435 |
| HG-U133 (A) | Agilent (E) | 12514 | 19514 | 10907 | 21121 | 0.516 |
| HG-U133 (A) | RNA-seq (F) | 12514 | 23808 | 12262 | 24060 | 0.510 |
| HG-U133 Plus 2.0 (B) | HG-U95 (C) | 20202 | 8590 | 8528 | 20264 | 0.421 |
| HG-U133 Plus 2.0 (B) | HuEx-1.0-ST (D) | 20202 | 28287 | 19905 | 28584 | 0.696 |
| HG-U133 Plus 2.0 (B) | Agilent (E) | 20202 | 19514 | 15838 | 23878 | 0.663 |
| HG-U133 Plus 2.0 (B) | RNA-seq (F) | 20202 | 23808 | 19284 | 24726 | 0.780 |
| HG-U95 (C) | HuEx-1.0-ST (D) | 8590 | 28287 | 8543 | 28334 | 0.302 |
| HG-U95 (C) | Agilent (E) | 8590 | 19514 | 7555 | 20549 | 0.368 |
| HG-U95 (C) | RNA-seq (F) | 8590 | 23808 | 8509 | 23889 | 0.356 |
| HuEx-1.0-ST (D) | Agilent (E) | 28287 | 19514 | 17555 | 30246 | 0.580 |
| HuEx-1.0-ST (D) | RNA-seq (F) | 28287 | 23808 | 22026 | 30069 | 0.733 |
| Agilent (E) | RNA-seq (F) | 19514 | 23808 | 16960 | 26362 | 0.643 |

Table S1: **Pairwise gene-set overlap across CellMiner platforms after gene-level collapsing.** For each ordered pair of platforms, $n_1$ and $n_2$ denote the number of unique Entrez gene IDs retained on Platform 1 and Platform 2 after probe-to-gene collapsing (median across probes, excluding Affymetrix `_x_at` probesets) and sample harmonization. $n_{\text{intersect}}$ and $n_{\text{union}}$ denote the size of the intersection and union of the two gene sets, respectively. The Jaccard index is computed as $n_{\text{intersect}}/n_{\text{union}}$ and summarizes cross-platform overlap in the available gene space used for model fitting and MSPE evaluation.

Since genes differ across platforms, to explore the sample-level agreement across platforms, we summarized expression per sample by median across genes within each platform, then compare those sample-level medians. By correlating these per-sample medians, we assessed the global similarity of the platforms. A heatmap (Figure S1A) of these spearman correlations shows low similarities between all platforms, so we expect bigger issues in downstream cross-platform comparisons. Density plots of pooled expression values (all values across common samples, Figure S1B) reveal that the platforms have vastly different dynamic ranges and scales. Within each platform, per-sample density overlays (Figure S1C) help spot outlier samples (e.g., one sample consistently shifted or unusually dispersed). Most curves overlap tightly within a platform, indicating normalization technique for each platform is likely consistent.

Across the six expression platforms, 57 matched samples were observed in common. We assessed cross-platform predictive accuracy using 10-fold cross-validation with a single set of fold assignments reused across all platforms and models. In each fold, PXN models and two MatchMixeR variants were trained on 90% of paired samples and evaluated on the held-out 10% by bidirectional prediction. A subset of 7,441 genes was identified as being measured across all six platforms. This common gene set serves as the primary basis for direct gene-level cross-platform comparisons. For each trained model and each ordered source-

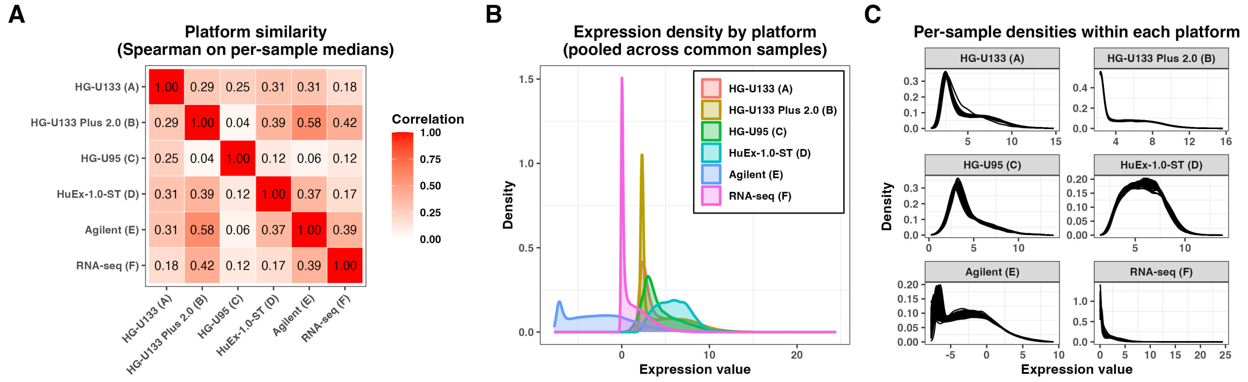


Figure S1: **Sample-level agreement across platforms for CellMiner Data.** (A) Heatmap of Spearman correlations of sample-level medians across genes across platforms. (B) Density plots of pooled expression values for all platforms(all values across common samples). (C) Per-sample density plots within each platform.

target platform pair it supports, we predicted target-platform expression from the source platform and computed the mean squared prediction error (MSPE) against the observed target expression in the same held-out samples. MSPE was computed on the gene set used to fit the corresponding model, ensuring that comparisons were made within a consistent feature space. For all six platforms, our evaluation produced a total of $\binom{6}{2} = 30$ ordered pairwise normalization tasks.

#### S6.B Data Preparation of TCGA Data

We conducted differential expression (DE) analysis using the TCGA dataset, comprising 14589 gene expression profiles from 582 paired samples—524 tumor and 58 normal samples — one pair of samples were excluded due to quality issues. Since the true DEGs are unknown, we defined a set of 6,602 DEGs based on the following procedure. First, we applied Limma [Smyth] separately to data from two different platforms, obtaining two lists of estimated tumor effects and corresponding p-values. We then identified *consistent genes* as those whose estimated tumor effects (in terms of log-fold changes, LFCs) were either both positive or both negative across platforms. From this subset, we selected DEGs according to two criteria: (a) an absolute estimated tumor effect of at least 0.3, and (b) a Benjamini-Hochberg (BH)-adjusted p-value less than 0.05. This resulting set of DEGs was treated as the gold standard.

It is worth noting that due to the substantial biological differences between tumor and normal tissues, almost all genes exhibited substantial LFCs that would significant inflation of type I error for the subsequent DE analysis. A histogram of the estimated LFCs for all genes is provided in Figure S2. To address this issue, we manually set the tumor-normal mean expression differences of non-DEGs (NDEGs) to zero for downstream DE analyses.

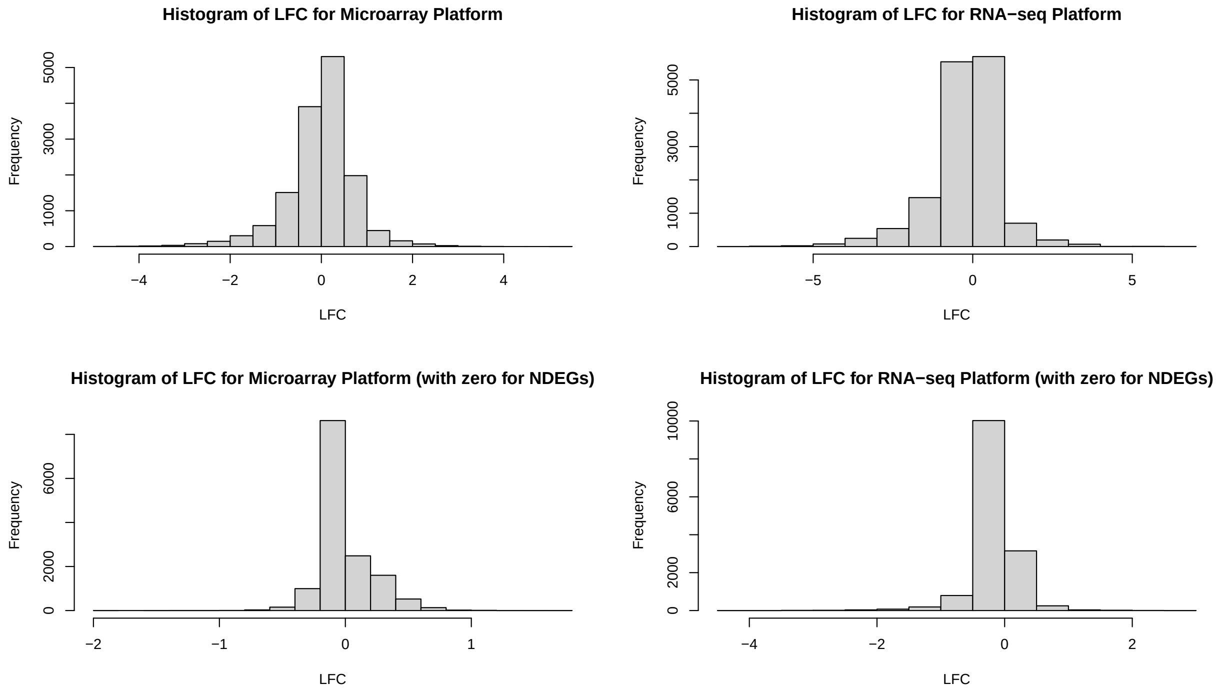


Figure S2: Histogram of the estimated LFCs for all genes in the TCGA data for real data demonstration.

### S7 Supplementary Tables for Real Data Demonstration

We additionally categorized applicable bridged models by whether the two component models share a platform ("platform overlap"), which determines whether the bridged model spans three platforms (overlap) or four platforms (disjoint) (Supplementary Table S2). Disjoint component pairs are common in both zero-shot and perturbed settings: among the 16 zero-shot bridged models, 12 are formed from disjoint pairs, and among the 14 perturbed models that include $[AB]$, 6 are also disjoint (Supplementary Table S2).

| Corresponding pairwise direct model $[AB]$ included | Two component models share a commom platform | Count | Example | | | |
|---|---|---|---|---|---|---|
| | | | Platforms for Component 1 | Platforms for Component 2 | Union Platforms | Shared Platforms |
| No (Zero-Shot) | Yes (overlap) | 4 | A,C | B,C | A,B,C | C |
| No (Zero-Shot) | No (disjoint) | 12 | A,C | B,D | A,B,C,D | – |
| Yes (Perturbed) | Yes (overlap) | 8 | A,B | B,D | A,B,D | B |
| Yes (Perturbed) | No (disjoint) | 6 | A,B | C,D | A,B,C,D | – |

Table S2: **Taxonomy of applicable bridged models for a representative directional task.** For the representative task $A \to B$, we classify the 30 *applicable* two-way bridged models (i.e., bridged models whose platform union contains both $A$ and $B$) by (i) whether the corresponding direct pairwise model $[AB]$ is included as a component (Yes: *perturbed*; No: *zero-shot*) and (ii) whether the two component pairwise models share a platform (overlap) or are disjoint. Overlap implies the bridged model spans three platforms, whereas disjoint component pairs span four platforms. The table reports counts in each category and provides a concrete example of the two component pairs, their union, and any shared platform.

| | **Zero-shot ($n = 16$) vs Perturbed ($n = 14$)** | | | |
|---|---|---|---|---|
| **Direction** | **MSPE$_{\text{zero}}$** | **MSPE$_{\text{perturb}}$** | **MSPE$_{\text{direct}}$** | $n_{\text{perturb-win}}$ |
| A → B | 0.5399 | 0.5277 | 0.5250 | 5 |
| B → A | 0.3723 | 0.3643 | 0.3598 | 1 |
| A → C | 0.1334 | 0.1234 | 0.1189 | 0 |
| C → A | 0.2522 | 0.2355 | 0.2293 | 2 |
| A → D | 0.2206 | 0.2116 | 0.2092 | 4 |
| D → A | 0.3055 | 0.2972 | 0.2917 | 0 |
| A → E | 0.8720 | 0.8304 | 0.8150 | 1 |
| E → A | 0.3672 | 0.3566 | 0.3524 | 2 |
| B → C | 0.2366 | 0.2336 | 0.2312 | 1 |
| C → B | 0.6126 | 0.5997 | 0.6016 | 7 |
| B → D | 0.1985 | 0.1910 | 0.1849 | 0 |
| D → B | 0.4499 | 0.4331 | 0.4237 | 0 |
| B → E | 0.7123 | 0.6898 | 0.6704 | 1 |
| E → B | 0.4819 | 0.4632 | 0.4532 | 0 |
| C → D | 0.2443 | 0.2359 | 0.2369 | 8 |
| D → C | 0.1868 | 0.1823 | 0.1791 | 1 |
| C → E | 0.9301 | 0.8928 | 0.8878 | 5 |
| E → C | 0.2276 | 0.2213 | 0.2189 | 2 |
| D → E | 0.5228 | 0.4896 | 0.4670 | 0 |
| E → D | 0.1569 | 0.1436 | 0.1352 | 0 |
| A → F | 0.4271 | 0.4234 | 0.4287 | 10 |
| F → A | 0.4000 | 0.3944 | 0.4167 | 11 |
| B → F | 0.4198 | 0.4120 | 0.4149 | 12 |
| F → B | 0.5537 | 0.5413 | 0.5550 | 8 |
| C → F | 0.4256 | 0.4243 | 0.4301 | 11 |
| F → C | 0.2457 | 0.2434 | 0.2507 | 12 |
| D → F | 0.3761 | 0.3618 | 0.3617 | 8 |
| F → D | 0.1997 | 0.1908 | 0.1940 | 10 |
| E → F | 0.3910 | 0.3772 | 0.3765 | 6 |
| F → E | 0.6888 | 0.6750 | 0.6835 | 8 |

Table S3: **Direction-level task summaries for zero-shot and perturbed bridging (model-averaged within each task).** For each ordered platform-to-platform normalization task (direction), we enumerate all *applicable* two-way bridged models whose platform union contains both the source and target platforms and compute within-task averages of MSPE over applicable models. MSPE$_{\text{zero}}$ is the mean MSPE across applicable *zero-shot* bridged models that do *not* include the corresponding direct source–target pairwise model as a component (n=16 per task). MSPE$_{\text{perturb}}$ is the mean MSPE across applicable *perturbed* bridged models that *do* include the corresponding direct pairwise model as a component but additionally bridge in extra platform structure (n=14 per task). MSPE$_{\text{direct}}$ is the MSPE of the corresponding direct pairwise PXN model for that direction. $n_{\text{perturb-win}}$ denotes the number of perturbed bridged models (out of 14) whose MSPE is lower than the direct baseline MSPE$_{\text{direct}}$ for that task.

| | Zero-shot, Disjoint ($n = 12$) | | Zero-shot, Overlap ($n = 4$) | | Perturbed, Disjoint ($n = 6$) | | | Perturbed, Overlap ($n = 8$) | | |
|---|---|---|---|---|---|---|---|---|---|---|
| **Direction** | $\text{MSPE}_{\text{zero}}$ | $\text{MSPE}_{\text{direct}}$ | $\text{MSPE}_{\text{zero}}$ | $\text{MSPE}_{\text{direct}}$ | $n_{\text{perturb-win}}$ | $\text{MSPE}_{\text{perturb}}$ | $\text{MSPE}_{\text{direct}}$ | $n_{\text{perturb-win}}$ | $\text{MSPE}_{\text{perturb}}$ | $\text{MSPE}_{\text{direct}}$ |
| A → B | 0.5396 | – | 0.5410 | – | 1 | 0.5371 | 0.5307 | 4 | 0.5207 | 0.5207 |
| B → A | 0.3732 | – | 0.3697 | – | 0 | 0.3719 | 0.3645 | 1 | 0.3586 | 0.3563 |
| A → C | 0.1325 | – | 0.1361 | – | 0 | 0.1255 | 0.1193 | 0 | 0.1219 | 0.1186 |
| C → A | 0.2512 | – | 0.2551 | – | 0 | 0.2400 | 0.2306 | 2 | 0.2321 | 0.2284 |
| A → D | 0.2201 | – | 0.2221 | – | 0 | 0.2143 | 0.2107 | 4 | 0.2096 | 0.2081 |
| D → A | 0.3058 | – | 0.3048 | – | 0 | 0.3034 | 0.2957 | 0 | 0.2925 | 0.2887 |
| A → E | 0.8706 | – | 0.8765 | – | 0 | 0.8352 | 0.8160 | 1 | 0.8268 | 0.8143 |
| E → A | 0.3687 | – | 0.3628 | – | 0 | 0.3652 | 0.3572 | 2 | 0.3502 | 0.3488 |
| B → C | 0.2362 | – | 0.2376 | – | 0 | 0.2361 | 0.2320 | 1 | 0.2317 | 0.2306 |
| C → B | 0.6128 | – | 0.6119 | – | 3 | 0.6059 | 0.6068 | 4 | 0.5950 | 0.5976 |
| B → D | 0.1976 | – | 0.2010 | – | 0 | 0.1952 | 0.1870 | 0 | 0.1879 | 0.1834 |
| D → B | 0.4530 | – | 0.4405 | – | 0 | 0.4498 | 0.4367 | 0 | 0.4206 | 0.4139 |
| B → E | 0.7054 | – | 0.7328 | – | 0 | 0.6939 | 0.6672 | 1 | 0.6868 | 0.6727 |
| E → B | 0.4849 | – | 0.4727 | – | 0 | 0.4788 | 0.4663 | 0 | 0.4514 | 0.4433 |
| C → D | 0.2441 | – | 0.2449 | – | 3 | 0.2376 | 0.2383 | 5 | 0.2346 | 0.2359 |
| D → C | 0.1866 | – | 0.1875 | – | 0 | 0.1853 | 0.1806 | 1 | 0.1800 | 0.1780 |
| C → E | 0.9288 | – | 0.9340 | – | 3 | 0.8928 | 0.8877 | 2 | 0.8928 | 0.8879 |
| E → C | 0.2281 | – | 0.2258 | – | 0 | 0.2257 | 0.2210 | 2 | 0.2180 | 0.2173 |
| D → E | 0.5154 | – | 0.5451 | – | 0 | 0.4895 | 0.4598 | 0 | 0.4897 | 0.4724 |
| E → D | 0.1566 | – | 0.1578 | – | 0 | 0.1480 | 0.1370 | 0 | 0.1402 | 0.1339 |
| A → F | 0.4270 | – | 0.4275 | – | 6 | 0.4244 | 0.4326 | 4 | 0.4226 | 0.4257 |
| F → A | 0.4007 | – | 0.3979 | – | 6 | 0.3996 | 0.4345 | 5 | 0.3905 | 0.4033 |
| B → F | 0.4243 | – | 0.4065 | – | 4 | 0.4262 | 0.4292 | 8 | 0.4014 | 0.4042 |
| F → B | 0.5587 | – | 0.5386 | – | 3 | 0.5621 | 0.5723 | 5 | 0.5257 | 0.5421 |
| C → F | 0.4249 | – | 0.4278 | – | 6 | 0.4237 | 0.4326 | 5 | 0.4247 | 0.4282 |
| F → C | 0.2457 | – | 0.2459 | – | 6 | 0.2451 | 0.2561 | 6 | 0.2421 | 0.2467 |
| D → F | 0.3810 | – | 0.3613 | – | 4 | 0.3783 | 0.3787 | 4 | 0.3495 | 0.3490 |
| F → D | 0.2002 | – | 0.1979 | – | 5 | 0.1964 | 0.2007 | 5 | 0.1866 | 0.1889 |
| E → F | 0.3968 | – | 0.3736 | – | 4 | 0.3938 | 0.3958 | 2 | 0.3647 | 0.3619 |
| F → E | 0.6840 | – | 0.7033 | – | 4 | 0.6788 | 0.6877 | 4 | 0.6722 | 0.6803 |
| **Average** | **0.4052** | – | **0.4047** | – | | **0.3987** | **0.3955** | | **0.3874** | **0.3860** |

Table S4: **Direction-level MSPE summaries stratified by zero-shot versus perturbed bridging and by overlap versus disjoint construction as explained in Table S2.** For each ordered normalization task, we summarize MSPE for *applicable* two-way bridged models stratified by whether the bridged model excludes the corresponding direct pairwise component (Zero-shot) or includes it (Perturbed), and by whether the two component pairwise models share a platform (Overlap) or not (Disjoint). $\text{MSPE}_{\text{zero}}$ and $\text{MSPE}_{\text{perturb}}$ denote the average MSPE across bridged models in the corresponding category. $\text{MSPE}_{\text{direct}}$ is the MSPE of the corresponding direct pairwise PXN model for that task (reported only for perturbed categories where the direct pairwise component is included). $n_{\text{perturb-win}}$ is the number of *perturbed* bridged models in the category whose MSPE is lower than $\text{MSPE}_{\text{direct}}$ for that task.

### References

G. K. Smyth. Limma: linear models for microarray data. In R. Gentleman, V. Carey, S. Dudoit, R. Irizarry, and W. Huber, editors, *Bioinformatics and Computational Biology Solutions Using R and Bioconductor*, pages 397–420. Springer.

S. Zhang, J. Shao, D. Yu, X. Qiu, and J. Zhang. MatchMixeR: a cross-platform normalization method for gene expression data integration. *Bioinformatics*, 36(8):2486–2491, Apr. 2020.
